# Neural Onset and Offset Responses in Core and Belt Auditory Cortex of Young and Aged Alert Macaque Monkeys

**DOI:** 10.1101/2025.05.29.656850

**Authors:** Danyi Lu, Gregg H. Recanzone

## Abstract

Age-related hearing loss is a ubiquitous malady in the geriatric population, yet the underlying neural mechanisms remain incompletely understood. We compared onset and offset responses of single neurons in the primary (A1), rostral (R), and caudolateral (CL) areas of the auditory cortex using spectral, spatial, and temporal stimuli in a young macaque monkey with normal-hearing and an aged macaque monkey with high frequency hearing loss. In the young monkey, core areas exhibited a significantly higher proportion of neurons responsive to both the onset and the offset of stimuli compared to the belt area CL, indicating functional differentiation. However, this distinction diminished with age, with the proportion of neurons with both onset and offset responses becoming more uniform across cortical areas. Onset firing rates were generally higher in the aged monkey, but with a lower signal-to-noise ratio, suggesting increased neural excitability but reduced response fidelity. Notably, CL neurons in the aged monkey exhibited a significantly greater disparity between the best frequencies for onset and offset responses, suggesting reduced spectral precision. Additionally, spectral tuning bandwidths (BW) were broader in the CL neurons in the aged monkey, while in the young monkey, A1 neurons exhibited significantly narrower onset BWs compared to offset BWs, a distinction that was lost in the aged monkey. These findings highlight a fundamental asymmetry in auditory cortical processing and suggest that belt area CL is particularly susceptible to age-related changes. Understanding these neural mechanisms provides insights into auditory aging and potential strategies for mitigating hearing deficits.

**New and Noteworthy:** This study finds that there are significant differences in the temporal fidelity of auditory cortical neuronal responses as a function of natural aging, particularly in the belt cortical field CL compared to core areas A1 and R. These differences are stimulus dependent and are consistent with known auditory processing deficits in geriatric human subjects.

## Introduction

Age-related hearing loss (ARHL), also known as presbycusis, is a complex disorder resulting from the gradual deterioration of auditory system processing as a consequence of natural aging. ARHL typically manifests as bilateral, symmetrical sensorineural hearing loss, most pronounced at higher frequencies (Gordon-Salant and Fitzgibbons, 1993; Frisina and Frisina, 1997; Recanzone, 2018; Yang et al., 2023). This progressive hearing impairment leads to social isolation and is associated with various comorbidities (WHO 2021). Additionally, increasing evidence links ARHL to cognitive decline and an elevated risk of dementia (Livingston et al., 2024). Despite ongoing research, critical gaps remain in understanding ARHL, particularly regarding how aging affects fundamental auditory processing mechanisms. Addressing these gaps is crucial for developing targeted interventions to mitigate the broader health consequences of ARHL and to improve the quality of life for aging populations.

Our laboratory has used the macaque monkey to study not only the peripheral deficits (Engle et al., 2013) but also both the electrophysiological (Ng et al., 2015) and histochemical changes (Gray, Engle et al., 2013; Gray, Rudolph et al., 2013; Engle et al., 2014; Gray et al., 2014a; Gray et al., 2014b) throughout the sub-cortical auditory neuroaxis as a function of aging. While we have observed changes consistent with those in both rodents and humans (see Gray and Recanzone, 2017), we have also shown that ARHL involves changes in cortical processing (Juarez-Salinas et al., 2010; Engle et al., 2014; see Recanzone, 2018; Ramamurthy and Recanzone, 2020), which plays a critical role in interpreting complex acoustic information. Critical stimulus features in auditory processing are transient events such as the onset and offset of stimuli. These transient events are commonly encoded as onset responses that occur within a few milliseconds of the onset of a stimulus, as well as offset responses that occur within a few milliseconds of the offset of a stimulus. Processing these transient events is crucial for auditory scene analysis, speech comprehension, and other higher-order auditory perceptions and functions as they help to parse the ongoing acoustic stream into discrete events. While extensive research has characterized onset responses in the auditory cortex, offset responses remain less well understood, particularly in primates.

Previous nonhuman primate studies have demonstrated that onset and offset responses of the neurons in the auditory cortex of alert monkeys exhibit distinct tuning properties in young animals, including differences in both spectral and spatial domains (Recanzone, 2000; Tian et al., 2013; Ramamurthy and Recanzone, 2017). When comparing primary auditory cortex (A1) and the caudolateral field (CL) in the auditory cortical belt (Kaas and Hackett, 2000), Ramamurthy and Recanzone (2017) found that while many neurons displayed overlapping onset and offset tuning profiles, a substantial proportion exhibited highly dissimilar tuning, particularly in the spectral domain. Building on these findings, they further investigated how aging affects the intensity coding of onset and offset responses (Ramamurthy and Recanzone, 2020). They reported that in aged monkeys without hearing loss, neurons exhibited higher firing rates and a shift toward more monotonic rate-level functions compared to young monkeys. Moreover, the similarity between onset and offset responses increased with aging, particularly in area CL.

Additionally, the impact of aging on the temporal processing capabilities of A1 neurons have also been explored (Overton and Recanzone, 2016; Ng and Recanzone, 2018). It was observed that neurons in A1 of the aged macaques with normal hearing exhibited increased spontaneous and driven firing rate, reduced phase-locking to amplitude-modulated (AM) stimuli, and a breakdown of the relationship between rate and temporal coding. These findings are consistent with the idea that aging alters the balance of excitation and inhibition in auditory cortical processing (Ouda et al., 2015; Gray and Recanzone, 2017; Recanzone, 2018), leading to changes in onset and offset responses. Given that speech perception relies on precise encoding of sound information (Gordon-Salant and Fitzgibbons, 1993; Frisina and Frisina, 1997), age-related changes in onset and offset response properties may underlie auditory processing deficits in the aging population (Qin et al., 2007; Kopp-Scheinpflug et al., 2018; Li et al., 2021).

Despite these advances, some key questions remain unanswered. Previous studies have compared neural responses between young monkeys and aged monkeys with normal hearing, revealing age-related differences in neuronal firing patterns. However, the neural responses of hearing-impaired aged monkeys remain unexplored, leaving a critical gap in our understanding of the cortical contributions to age-related hearing deficits. Furthermore, while previous research has separately characterized onset and offset tuning properties and examined the effects of aging on intensity coding, a comprehensive comparison of onset and offset responses in A1 and CL across young and aged monkeys, incorporating multiple stimulus dimensions, is still lacking. Specifically, how aging influences the spectral, spatial, and temporal dynamics of onset and offset responses remains an open question. Finally, it is unclear whether the observed changes in onset and offset tuning are specific to A1 and CL or extend to other auditory cortical fields.

To address these gaps, this study systematically investigates onset and offset responses in a young monkey with normal hearing and an aged monkey with high frequency hearing loss, providing a direct comparison of how natural aging and the common consequence of high-frequency hearing loss affects auditory cortical processing. Unlike previous studies that focused primarily on A1 and CL, we extend our analysis to include the rostral field (R), within the core of auditory cortex, enabling a broader assessment of onset and offset processing across core (area A1 and area R) and belt areas (area CL). We examine changes between these two macaques along three acoustic dimensions: spectral, spatial, and temporal, focusing on the proportion of cells exhibiting significant responses to both the onset and offset of these stimuli, the distinctions in the firing rates of onset and offset responses, and tuning properties of these neurons. This comprehensive approach enhances our understanding of the cortical consequences of natural aging.

## Materials and Methods

### 1. Ethical statement

All procedures used in these experiments were in accordance with AAALAC International and NIH guidelines for the use of animals in research and were approved by the Institutional Animal Care and Use Committee of the University of California, Davis.

### 2. Animals

The experiments were performed on two adult male rhesus macaque monkeys, aged 12-14 years (Monkey Y) and 26-27 years (Monkey O). Macaque monkeys are generally thought to age at approximately three times the rate of humans (Davis and Leathers, 1985), making Monkey Y the equivalent of 36-42 human years old and Monkey O the equivalent of 78-81 human years old at the time of data collection. Both animals had no history of ear infections or exposure to ototoxic drugs or excessive loud noise. Monkey Y had normal behavioral audiograms for tones from 500 to 16,000 Hz, while Monkey O was identified as having high frequency (>8,000 Hz) hearing loss (threshold > 70 dB SPL) based on ABR recordings (Ng et al., 2015).

### 3. Acoustic stimulus presentation

Experiments were conducted in a double-walled anechoic chamber (IAC) lined with sound-absorbent foam (Sonex 4-inch). During recording sessions, each monkey sat in an acoustically transparent primate chair with its head restrained at the center of a circular array. This array was 1.8 m in diameter with 16 speakers equally spaced 22.5° apart, spanning a full 360° in azimuth at ear level. Each sound was presented from one of these 16 locations during the experiment. Subjects passively listened to various auditory stimuli and were given diluted fruit juices at random intervals to maintain alertness.

Acoustic stimuli were generated using System 2 hardware (Tucker-Davis Technologies). Twelve repetitions of each stimulus were presented in a randomly interleaved fashion. This report describes the results from three different stimulus types which we refer to as ‘experiments’. In the spectral experiment (TONE), pure tone stimuli with a duration of 200 ms (5 ms rise/fall) were presented at 50 dB sound pressure level (SPL) from the speaker located directly opposite the ear (+90 deg) contralateral to the recorded hemisphere. This experiment included 14 tones spanning approximately 6 octaves in ½ octave steps (in kHz: 0.25, 0.353, 0.5, 0.707, 1.0, 1.41, 2.0, 2.83, 4.0, 5.66, 8.0, 11.31, 16.0, 19.60). In the spatial experiment (SP), noise-burst stimuli consisting of 200 ms duration (5 ms rise/fall) “unfrozen” Gaussian noise were presented at 60 dB SPL from locations spaced at 22.5 degrees across 360 degrees in azimuth at 0 degrees elevation. In the temporal experiment (AM), stimuli were 100% sinusoidally amplitude-modulated Gaussian noise of 500 ms duration. A sequence of 8 modulation frequencies (4, 8, 16, 32, 64, 128, 256, and 512 Hz) was presented at 60 dB SPL from directly opposite the contralateral ear (+90 deg).

### 4. Recording procedures

Monkeys were implanted with a restraining head post and a recording cylinder, enabling orthogonal penetration of the superior surface of the superior temporal gyrus in the left hemisphere (Recanzone et al., 2000a; Overton et al., 2017). A tungsten electrode (FHC) was inserted through a guide tube and lowered with a microdrive (Thomas Recording) during recording sessions. Neuronal activity was displayed on an oscilloscope and via a speaker outside the sound isolation chamber. Extracellular neural signals were passed through an amplifier (A-M Systems model 1700) and band-pass filtered at 0.3-10 kHz before being sent to an analog-to-digital converter (CED Power 1401), sampled at a rate of 50 kHz, and saved to a hard disk. The electrode was advanced while search stimuli were presented (broadband noise, tones, bandpass noise, clicks, etc.) to find acoustic-driven activity. Once auditory responses were encountered the multiple unit response, consisting of 1-5 single units, were manually characterized to define the threshold, characteristic frequency and responses to tones and noises of different bandwidths. Action potentials (spikes) were sorted offline using Spike2 (CED). The assignment of each recording site to areas R, A1, or CL was based on its location within the recording cylinder and its physiological characteristics (Recanzone et al., 2000a).

### 5. Data analysis

#### Firing rate analysis

Data collection and analysis were performed using custom scripts in MATLAB (MathWorks). Peristimulus time histograms (PSTHs) were constructed by counting the number of spikes that fell within each 1-ms bin. A 5-ms moving average window was applied to PSTHs solely for the visualization purpose in the figures. Unsmoothed PSTHs were used for analysis.

The spontaneous rate was calculated over the 100 ms pre-stimulus period, averaged across all stimulus presentations, and used as the baseline for subsequent analysis. Onset and offset responses were measured within 100 ms following stimulus onset and offset, respectively. Reponses were considered significant if the firing rate within the time window was either greater than or less than the baseline firing rate by more than two standard deviations (Recanzone et al., 2000a, b). The stimulus that evoked the highest firing rate was defined as the best frequency in kHz (for TONE or AM) or best location in degrees (for SP) of the neuron, corresponding to the onset and offset responses, respectively.

For comparisons between cells across different cortical areas and monkeys, normalization was applied on a cell-by-cell basis. We first calculated the average spontaneous rate measured across all of the stimuli for that cell as noted above. The onset and offset firing rates were then divided by that value.

#### Bandwidth of spectral, spatial, and temporal tuning curve

The bandwidth (BW) for spectral, spatial, and temporal tuning was defined as the width at half the maximum firing rate of the respective tuning curve. Firing rates for each frequency or direction were used to construct the turning curve. To calculate the BW, the maximum value was identified, and the range exceeding half of this peak was determined. BWs were calculated separately for onset and offset responses, as well as for each type of stimulus. The BWs for TONE and AM stimuli were calculated in the number of octaves, and the BWs for SP stimuli were calculated by the range in degrees.

The BWs of each peak for the spectral and temporal tuning were calculated as:

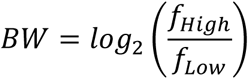

where *f*_*High*_ and *f*_*Low*_ are the two edges of the peak. For cases where 50% of best stimulus response occurs between two frequencies, the *f*_*High*_ and *f*_*Low*_ are obtained by taking the average value of these two frequencies.

The BWs of each peak in the spatial turning curve was calculated as:

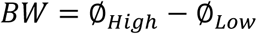

where ∅_*High*_ and ∅_*Low*_ are the degrees to the right and left of the peak. For cases where 50% of best stimulus response occurs between two speakers, the ∅_*High*_ and ∅_*Low*_ are obtained by taking the average value of these two degrees. If ∅_*High*_ is smaller than ∅_*Low*_ because the peak covers the 𝜋 direction, ∅_*High*_ is calculated as ∅_*High*_ + 2𝜋.

For multipeaked curves, the BWs were calculated by summing the BWs of each peak of the tuning curves.

#### Similarity of onset and offset responses

The similarity between onset and offset responses was calculated based on spectral, spatial, and temporal tuning curves for all neurons that exhibited both significant onset and offset responses. Best stimulus difference (BSD) was the absolute difference between the best onset stimulus and the best offset stimulus. The similarity index (SI) was used to represent BSD magnitude in octaves for TONE and AM stimuli, and in degrees for SP stimuli.

For spectral and temporal experiments:

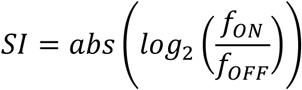

where *f*_*ON*_ and *f*_*OFF*_ are the best frequencies for the onset and offset responses, respectively.

For the spatial experiment:

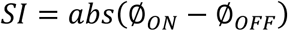

where ∅_*ON*_ and ∅_*OFF*_ are the best directions for the onset and offset responses, if *abs*(∅_*ON*_ − ∅_*OFF*_)> 𝜋, *SI* = 2𝜋 − *abs*(∅_*ON*_ − ∅_*OFF*_).

We report uncorrected p-values for all statistical tests exactly, so readers can draw their own conclusions (Heroux 2016).

## Results

### 1. Differences in the proportion of ON+OFF neurons

We recorded the responses of single auditory cortical neurons in alert macaque monkeys to three different types of acoustic stimuli: spectrally-varying 200 ms tones (TONE), spatially-varying broadband noise across acoustic space in azimuth (SP), and temporally-varying 100% amplitude modulated broadband noise (AM). Example responses for three different neurons, one for each stimulus type, are shown in Figure 1. For each neuron one can see clear onset responses, with clear offset responses seen for the spatial task (Fig. 1B). The onset and offset responses were considered significant if they differed from the baseline by at least 2 SD, where the baseline was defined as the average of the spontaneous rate measured across all stimuli. A cell was classified as an ON cell if at least one stimulus evoked a significant onset response, and as an OFF cell if at least one stimulus evoked a significant offset response. Combining these criteria, cells were categorized as ON only, OFF only, or ON+OFF cells (Table 1). This report is based on recordings of 460 single neurons from the core (areas R and A1) and belt (area CL) auditory cortex in two awake monkeys. This database is made up of 249 neurons in A1, 107 neurons in area R and 104 neurons in area CL. Only ON+OFF neurons were included in the subsequent analyses.

**Figure 1.**
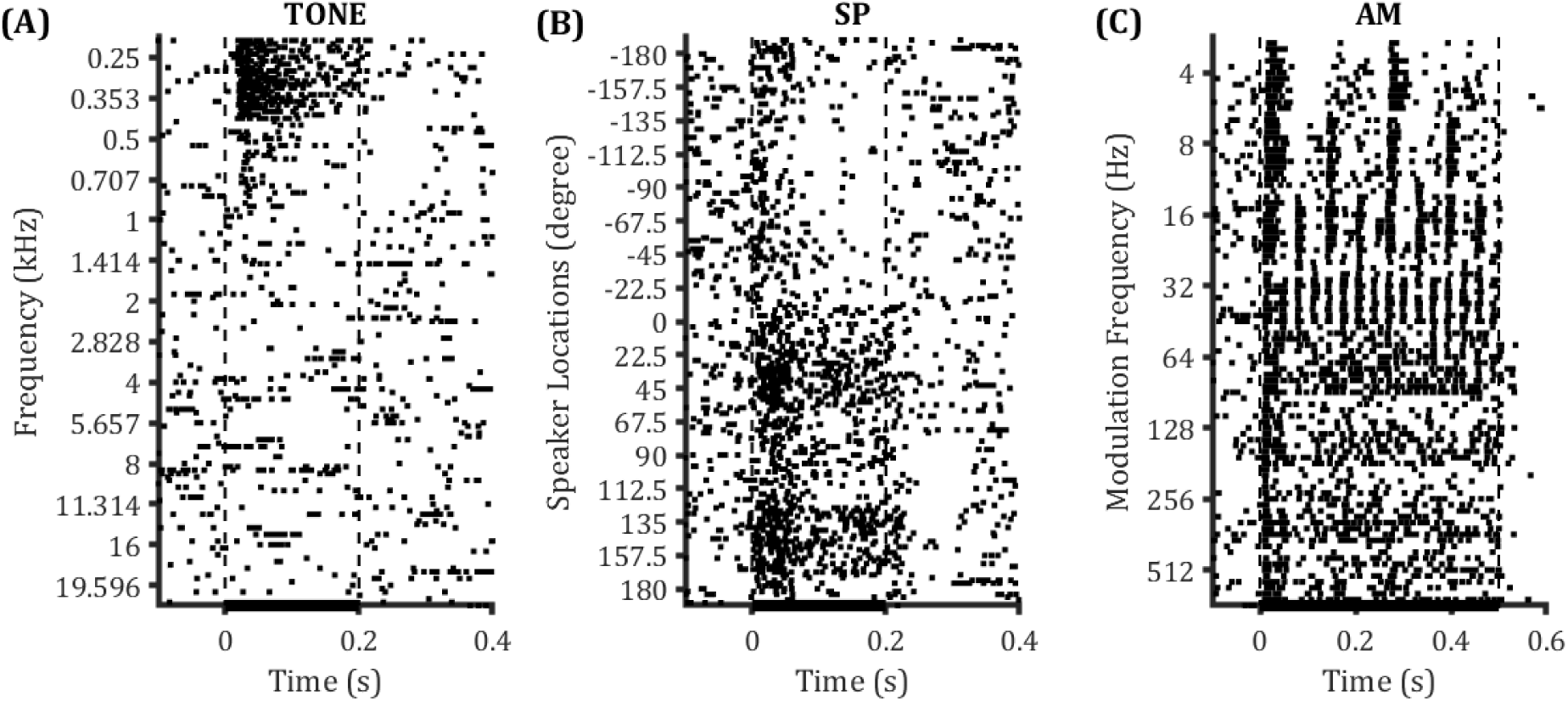
Raster plots from 3 representative A1 cells of monkey Y show good tuning for TONE (A), SP (B) and AM (C) stimuli, respectively. A: Tone frequency is shown on the y-axis, vertical lines show stimulus onset (left) and offset (right). This neuron had a robust ON response to low frequencies but little if any OFF response. B: Sptial location is shown on the y-axis. This cell had both ON and OFF responses in contralateral space. C: AM frequency is shown on the y-axis. There is good temporal precision for AM frequencies up to about 64 Hz, with a robust ON response across frequencies but a minimal OFF response.

**Table 1.** Percentages of R, A1, and CL neurons displaying onset and offset responses. Table 1. Total numbers of neurons exhibiting significant onset and offset responses are listed above. In the cell count column, the denominator represents the number of cells that showed either onset or offset responses greater than 2 SD for at least one stimulus. The numerator indicates, under each specific stimulus condition, the number of cells with onset or offset responses exceeding 2 SD. A cell was classified as ON only if at least one stimulus evoked a significant onset response without any significant offset response. Similarly, a cell was classified as OFF only if at least one stimulus induced a significant offset response with no significant onset response. ON+OFF cells refer to those that exhibited both significant onset and offset response to at least one stimulus.

|  | Monkey Y |  |  |  | Monkey O |  |  |  |
| --- | --- | --- | --- | --- | --- | --- | --- | --- |
|  | Cell Count | ON Only | OFF Only | ON+OFF | Cell Count | ON Only | OFF Only | ON+OFF |
| R-TONE | 88.6%<br>(39/44) | 20.5%<br>(8/39) | 15.4%<br>(6/39) | 64.1%<br>(25/39) | 61.9%<br>(39/63) | 17.9%<br>(7/39) | 7.7%<br>(3/39) | 74.4%<br>(29/39) |
| A1-TONE | 77.2%<br>(85/110) | 25.9%<br>(22/85) | 9.4%<br>(8/85) | 64.7%<br>(55/85) | 54.7%<br>(76/139) | 28.9%<br>(22/76) | 6.6%<br>(5/76) | 64.5%<br>(49/76) |
| CL-TONE | 54.1%<br>(33/61) | 24.2%<br>(8/33) | 15.2%<br>(5/33) | 60.6%<br>(20/33) | 51.2%<br>(22/43) | 31.8%<br>(7/22) | 9.1%<br>(2/22) | 59.1%<br>(13/22) |
| R-SP | 86.4%<br>(38/44) | 26.3%<br>(10/38) | 2.6%<br>(1/38) | 71.1%<br>(27/38) | 65.1%<br>(41/63) | 14.6%<br>(6/41) | 17.1%<br>(7/41) | 68.3%<br>(28/41) |
| A1-SP | 80.9%<br>(89/110) | 27.0%<br>(24/89) | 3.4%<br>(3/89) | 69.7%<br>(62/89) | 54.7%<br>(76/139) | 18.4%<br>(14/76) | 7.9%<br>(6/76) | 73.7%<br>(56/76) |
| CL-SP | 52.5%<br>(32/61) | 31.3%<br>(10/32) | 15.6%<br>(5/32) | 53.1%<br>(17/32) | 53.5%<br>(23/43) | 21.7%<br>(5/23) | 21.7%<br>(5/23) | 56.5%<br>(13/23) |
| R-AM | 77.3%<br>(34/44) | 29.4%<br>(10/34) | 2.9%<br>(1/34) | 67.6%<br>(23/34) | 57.1%<br>(36/63) | 16.7%<br>(6/36) | 25.0%<br>(9/36) | 58.3%<br>(21/36) |
| A1-AM | 80.9%<br>(89/110) | 34.8%<br>(31/89) | 9.0%<br>(8/89) | 56.2%<br>(50/89) | 53.2%<br>(74/139) | 25.7%<br>(19/74) | 9.5%<br>(7/74) | 64.9%<br>(48/74) |
| CL-AM | 52.5%<br>(32/61) | 53.1%<br>(17/32) | 12.5%<br>(4/32) | 34.4%<br>(11/32) | 51.2%<br>(22/43) | 18.2%<br>(4/22) | 22.7%<br>(5/22) | 59.1%<br>(13/22) |

As can be seen from Table 1, the vast majority of neurons were in some way responsive to at least one of these stimuli, and most commonly all three. It should be noted that all neurons encountered with an acoustically-driven response to any tested stimulus is revealed in the denominator of column 2 and 6 but not necessarily responsive to one of these three stimulus classes, for example those that were responsive only to narrow-band noise (see Rauschecker et al., 1995; Recanzone et al., 2000a; Rauschecker and Tian, 2004). The category of ON+OFF was prevalent across monkeys and cortical areas (columns 5 and 9).

Figure 2 shows the proportion of ON+OFF cells across three different areas in each monkey (left), and the same data are reproduced to compare these areas between the two monkeys (right). Chi-square tests were used across all comparisons and significance levels are indicated as p<0.01 (**), and p<0.05(*). Detailed statistical results are provided in Table In the young monkey, the proportion of ON+OFF neurons did not differ across the three types of stimuli between areas R and A1; however, the proportion of neurons in areas R and A1 was significantly higher than those in CL. This regional difference was not observed in the aged monkey. Additionally, comparing the two monkeys, the proportion of ON+OFF neurons in A1 was significantly higher in the young monkey in the TONE and SP experiments compared to the aged monkey, whereas no significant difference was found between area A1 of these two monkeys in the AM experiment, nor area R and area CL across all stimulus types. Thus, there were many similarities between monkeys, but we did note differences between the core (areas A1 and R) and belt (area CL) areas in the young monkey that were not apparent in the old monkey, due at least in part to differences in A1 between the old and young monkeys.

**Figure 2.**
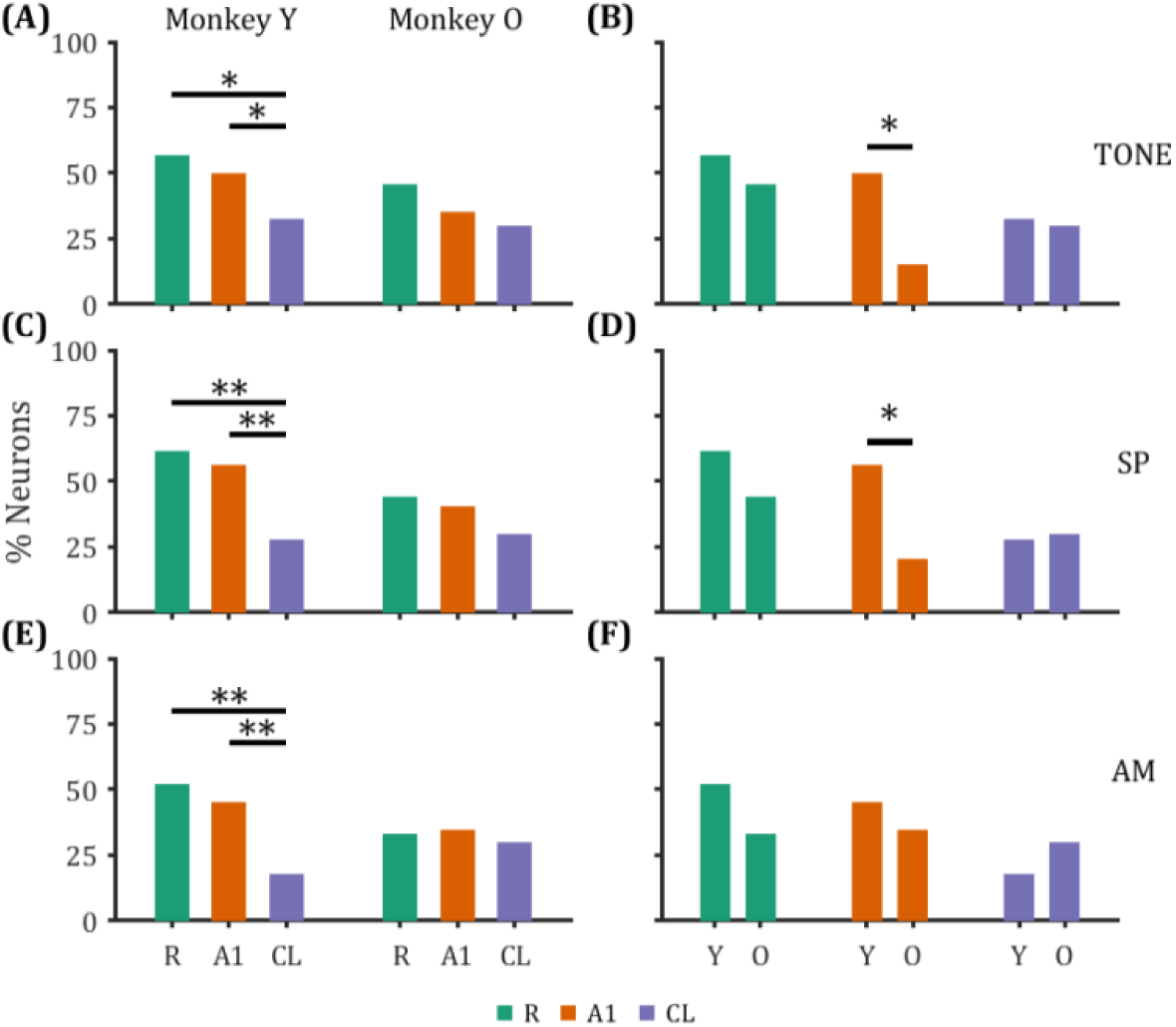
Comparison of the percentage of ON+OFF cells. A, C, E: comparison of the ON+OFF cells percentages within the young monkey (Monkey Y) and the aged monkey (Monkey O) across R, A1, and CL areas. B, D, F: comparison of the ON+OFF cells percentages between Monkey Y and Monkey O across R, A1, and CL areas. The percentages of significant neurons differed between core (A1 and R) and belt (CL) cortical areas in monkey Y but not monkey O (left). However, differences between monkeys were only seen in A1 for the TONE and SP experiments (**: p<0.01; *: p<0.05; chi-square test).

### 2. Differences in the best onset and best offset firing rates

After analyzing the differences in the proportion of ON+OFF cells, we aimed to further investigate how the firing patterns of these cells varied across different regions and between the two monkeys. First, we applied the Shapiro-Wilk test to assess the normality of the data. Upon finding that the data were non-parametric, we chose the two-sample Kolmogorov-Smirnov test to evaluate the differences between the datasets.

We used the same dataset to compare the firing rates between the different areas within each monkey (Fig. 3; statistical details in Table 3) for both the best onset responses (Fig. 3A-C) as well as offset responses (Fig. 3D-F). The results showed that in the TONE experiment, for both onset and offset responses, there was only one significant difference among the three areas within the two monkeys: in the aged monkey, area R, showed a higher firing rate than A1 for the offset responses. This difference reached the p < 0.05 statistical criteria (Fig. 3D, right panel; p = 0.0213, see Table 3). This contrasts with the spatially-varying SP experiment. For the onset responses in the young monkey, CL had the highest onset firing, followed by A1, with R showing the lowest firing rate (Fig. 3B, left panel). For the offset response, this pattern was reversed but only the difference between A1 and CL reached statistical significance. (Fig. 3E, left panel). The old monkey showed a different pattern of onset firing with firing rates for each cortical area significantly different than the other two (Fig. 3B, right panel). Interestingly, the offset firing rates showed the opposite pattern than the pattern of onset firing rates of the young monkey, also where significant differences were noted between each cortical area (Fig. 3E, right panel). Finally, the results from the AM experiment showed different patterns than those of the other two experiments for both onset and offset responses in both monkeys. The onset responses of R and A1 in the young monkey showed higher firing rates than CL (Fig. 3C, left panel). For all other comparisons, there was no significant difference in the offset responses in the young monkey, nor in the onset and offset responses in the aged monkey.

**Figure 3.**
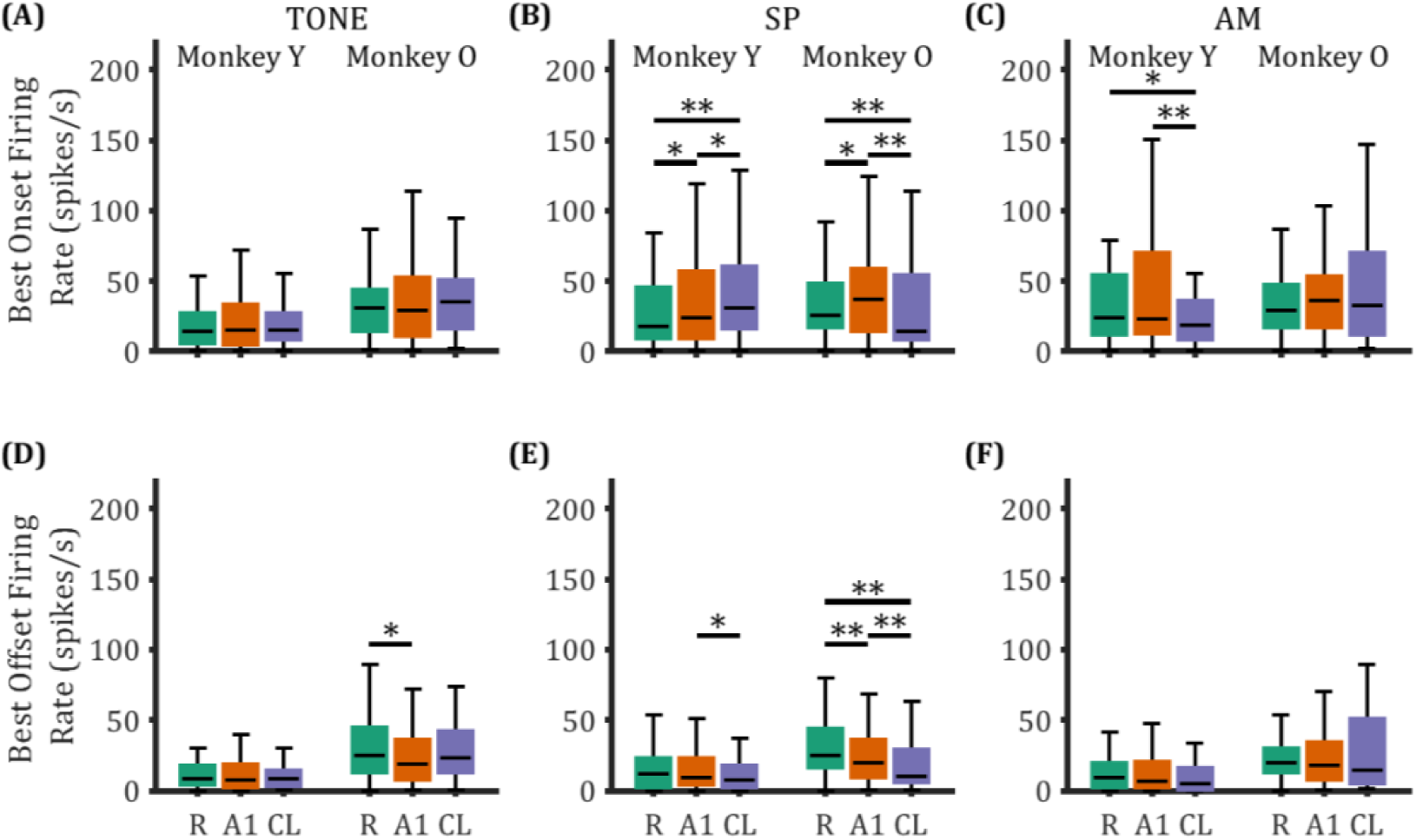
Comparison of the best onset and best offset firing rates across different areas of the auditory cortex in the young and aged monkeys. Box plots illustrating the onset (A-C) and offset (D-F) firing rates across three cortical areas between Monkey Y and Monkey O. The boxes represent the interquartile range from the 25th percentile to the 75th percentile, with the black horizontal bars indicate the median values. The whiskers extend from the edges of the box to the minimum and maximum values within 1.5 times the IQR from 25% and 75%, respectively. Any data points beyond the whiskers are considered outliers [**: p<0.01; *: p<0.05; two-sample Kolmogorov-Smirnov (K-S) test].

**Table 2.** Statistical details of the proportion of neurons. Table 2. Each cortical area (columns 3-5) shows the percentage of ON+OFF cells, for the young (Y) and old (O) monkeys (column 2) for each of the three stimulus types (rows). Statistical p-values are shown in columns 6 (between areas) and 7 (between monkeys), those that are considered statistically significant are highlighted in yellow when p<0.01 or in green when 0.01<p<0.05 (Chi-square test). There were no differences between cortical areas in the old monkey, but consistent differences between the core and belt areas in the young monkey for all three stimulus types.

|  | Monkey | R | A1 | CL | Among Areas | Between Monkeys |
| --- | --- | --- | --- | --- | --- | --- |
| ON+OFF cells % (TONE) | Y | 56.8% (25/44) | 50.0% (55/110) | 32.8% (20/61) | R≈A1: p=0.444<br>A1>CL: p=0.0298<br>R>CL: p=0.0141 | R: Y≈O, p=0.272<br>A1: Y>O, p=0.0191<br>CL: Y≈O, p=0.783 |
|  | O | 46.0% (29/63) | 35.3% (49/139) | 30.2% (13/43) | R≈A1: p=0.145<br>A1≈CL: p=0.544<br>R≈CL: p=0.102 |  |
| ON+OFF cells % (SP) | Y | 61.4% (27/44) | 56.4% (62/110) | 27.9% (17/61) | R≈A1: p=0.570<br>A1>CL: p=3.43e-4<br>R>CL: p=5.99e-4 | R: Y≈O, p=0.0849<br>A1: Y>O, p=0.0116<br>CL: Y≈O, p=0.0793 |
|  | O | 44.4% (28/63) | 40.3% (56/139) | 39.5% (17/43) | R≈A1: p=0.579<br>A1≈CL: p=0.235<br>R≈CL: p=0.140 |  |
| ON+OFF cells % (AM) | Y | 52.3% (23/44) | 45.5% (50/110) | 18.0% (11/61) | R≈A1: p=0.444<br>A1>CL: p=3.36e-4<br>R>CL: p=2.16e-4 | R: Y≈O, p=0.0501<br>A1: Y≈O, p=0.0798<br>CL: Y≈O, p=0.146 |
|  | O | 33.3% (21/63) | 34.5% (48/139) | 30.2% (13/43) | R≈A1: p=0.868<br>A1≈CL: p=0.602<br>R≈CL: p=0.737 |  |

**Table 3.**
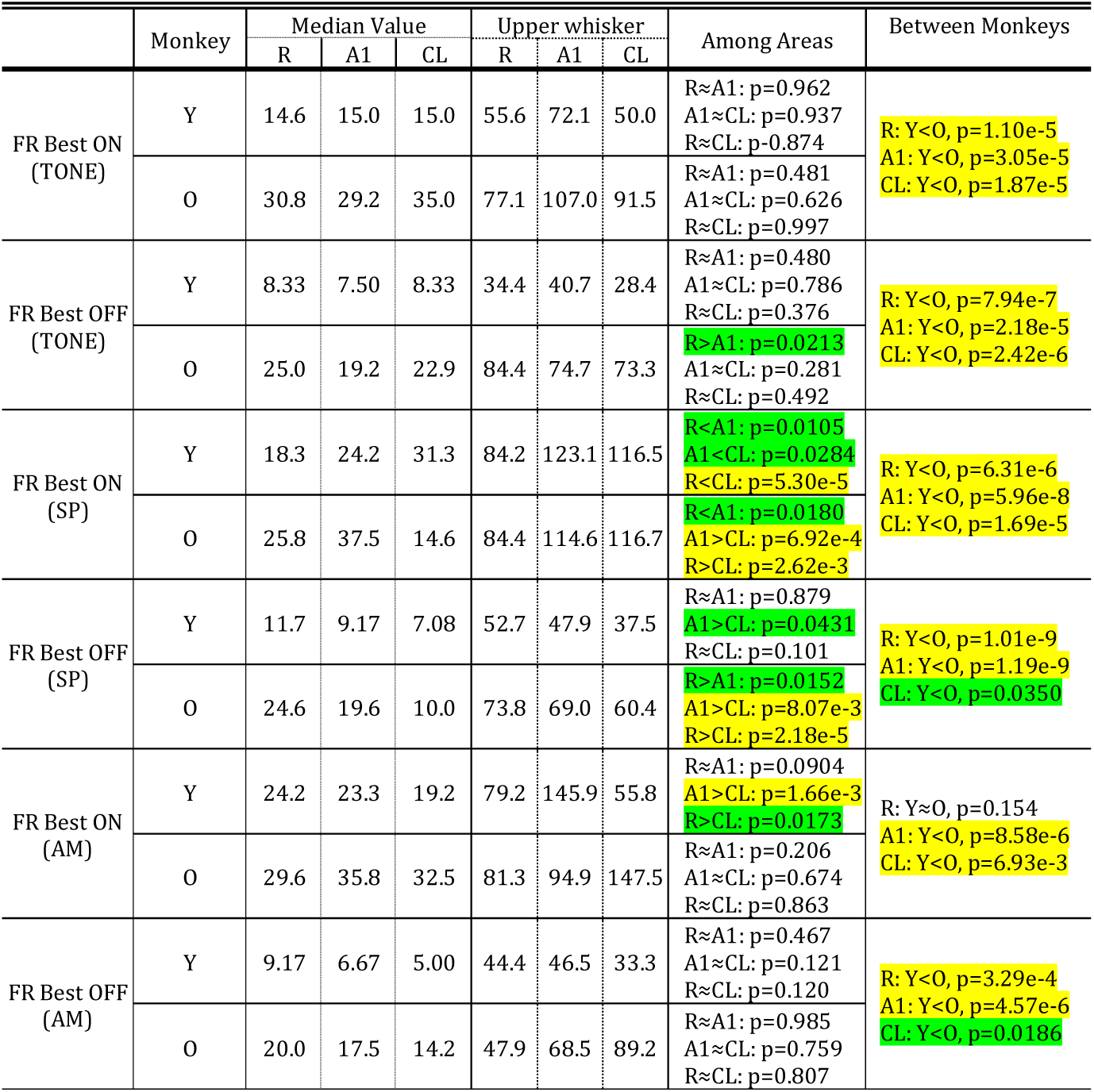
Statistical details from the analysis shown in Figures 3 and 4. Table 3. Each monkey (Y) and (O) are shown in alternating rows for each of the three stimulus types divided by rows into onset (upper) and offset (lower) rows. Median value of the firing rates corresponding to the median line. Statistical results with p<0.01 are highlighted in yellow and 0.01<p< 0.05 are in green (two-sample K-S test). There were consistent differences between the young and old monkeys across comparisons (far right column).

The preceding analysis compared the firing rates between cortical areas for each monkey individually. We next focused the analysis on examining potential differences in the magnitude of the best onset and best offset responses in various cortical areas and stimuli between the two monkeys (Fig. 4; statistical details in the far right column of Table 3). In the old monkey, most of the firing rates in the three different cortical areas in response to the three types of stimuli, whether during onset or offset, were significantly higher than those in the young monkey (right side of each pair of colored bars; Fig. 4). With two exceptions, one was in the SP experiment, the onset firing rate in CL neurons of the young monkey was higher than the aged monkey (Fig. 4B; purple bars); the other was in the AM experiment, where there was no significant difference in the onset response of neurons in R between the two monkeys (Fig. 4C; green bars). Thus, consistent with previous reports from different monkeys, the aged monkey in this study had a similarly elevated firing rate compared to the younger control (see Recanzone, 2018).

**Figure 4.**
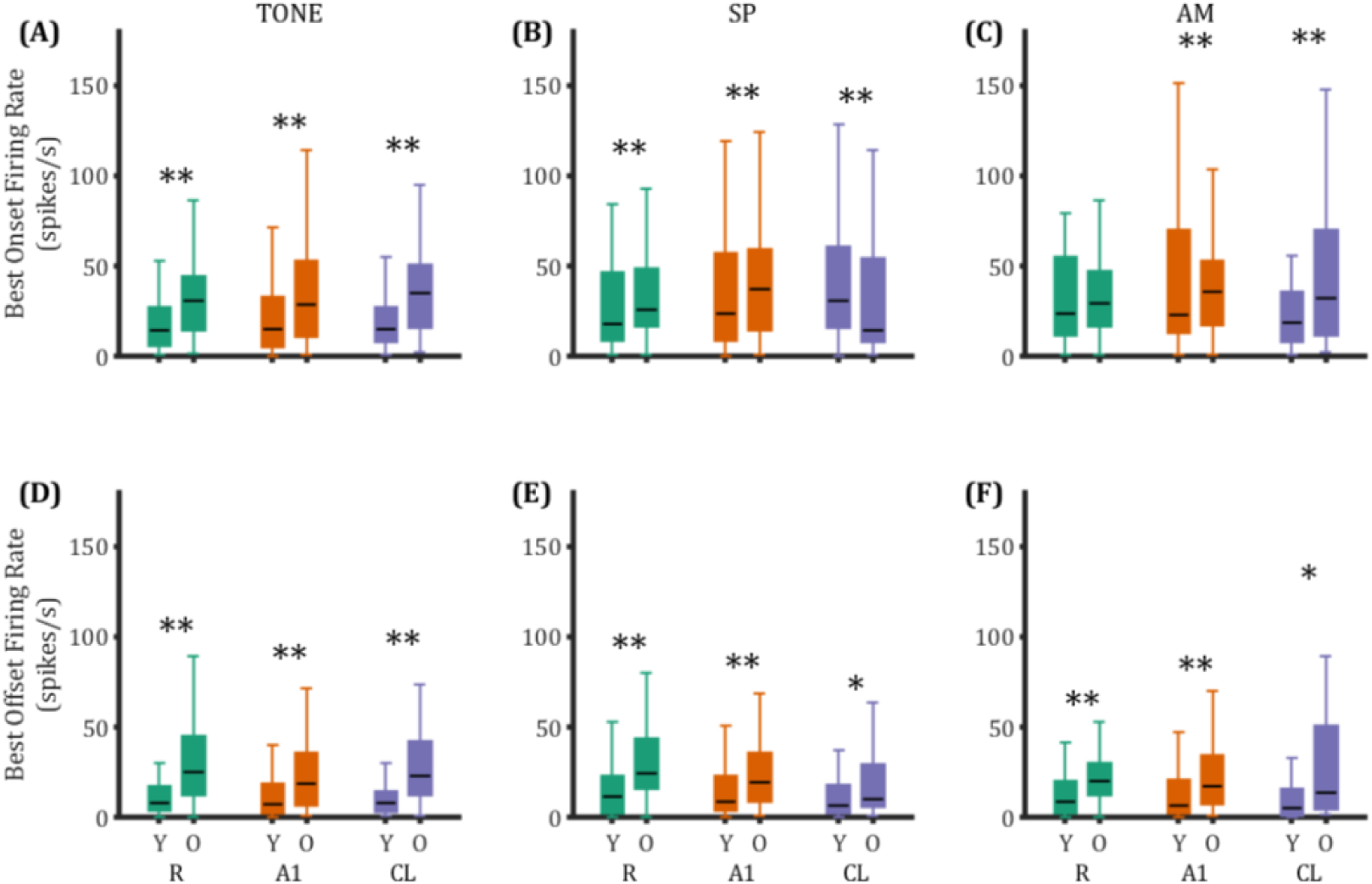
Comparison of the best onset and best offset firing rates between the young and aged monkeys. Box plots illustrating the onset (A-C) and offset (D-F) firing rates across three cortical areas between Monkey Y and Monkey O. The boxes represent the interquartile range from the 25th percentile to the 75th percentile, with the black horizontal bars indicate the median values. The whiskers extend from the edges of the box to the minimum and maximum values within 1.5 times the interquartile range (IQR) from 25% and 75%, respectively. Any data points beyond the whiskers are considered outliers (**: p<0.01; *: p<0.05; two-sample K-S test).

To continue the comparison between the two monkeys we also compared the firing rates normalized by the spontaneous activity (Fig. 5; detailed statistical results in Table 4), Previous studies have shown that the spontaneous activity of auditory cortical neurons is higher in aged compared to younger monkeys (see Recanzone, 2018), which also proved to be the case in these two different animals (area R, Monkey Y median = 5.83 < Monkey O median = 8.33, p=7.13e-12; area A1, Monkey Y median = 5.83 < Monkey O median = 10.3, p=1.75e-8; area CL, Monkey Y median = 4.17 < Monkey O median = 13.3, p=8.50e-7). While we report the median value for convenience, the K-S test used all values. For this study, we found that, in most cases, the young monkey exhibited a higher evoked firing rate than the old monkey at a statistically significant level once spontaneous activity was taken into account. In every case, the normalized firing rate for the young monkey was statistically significantly greater than that for the older monkey (Fig. 5A-C). For the offset responses, while the young monkey often had a greater normalized firing rate, the difference did not reach statistical significance in 4/9 comparisons (Fig. 5D-F) which may be due in part to the relatively low firing rates. Thus, it appears that while the overall firing rate was higher in the old monkey, the signal-to-noise ratio was lower for both onset and offset responses, similar to previous findings in old monkeys with normal hearing (see Recanzone, 2018).

**Figure 5.**
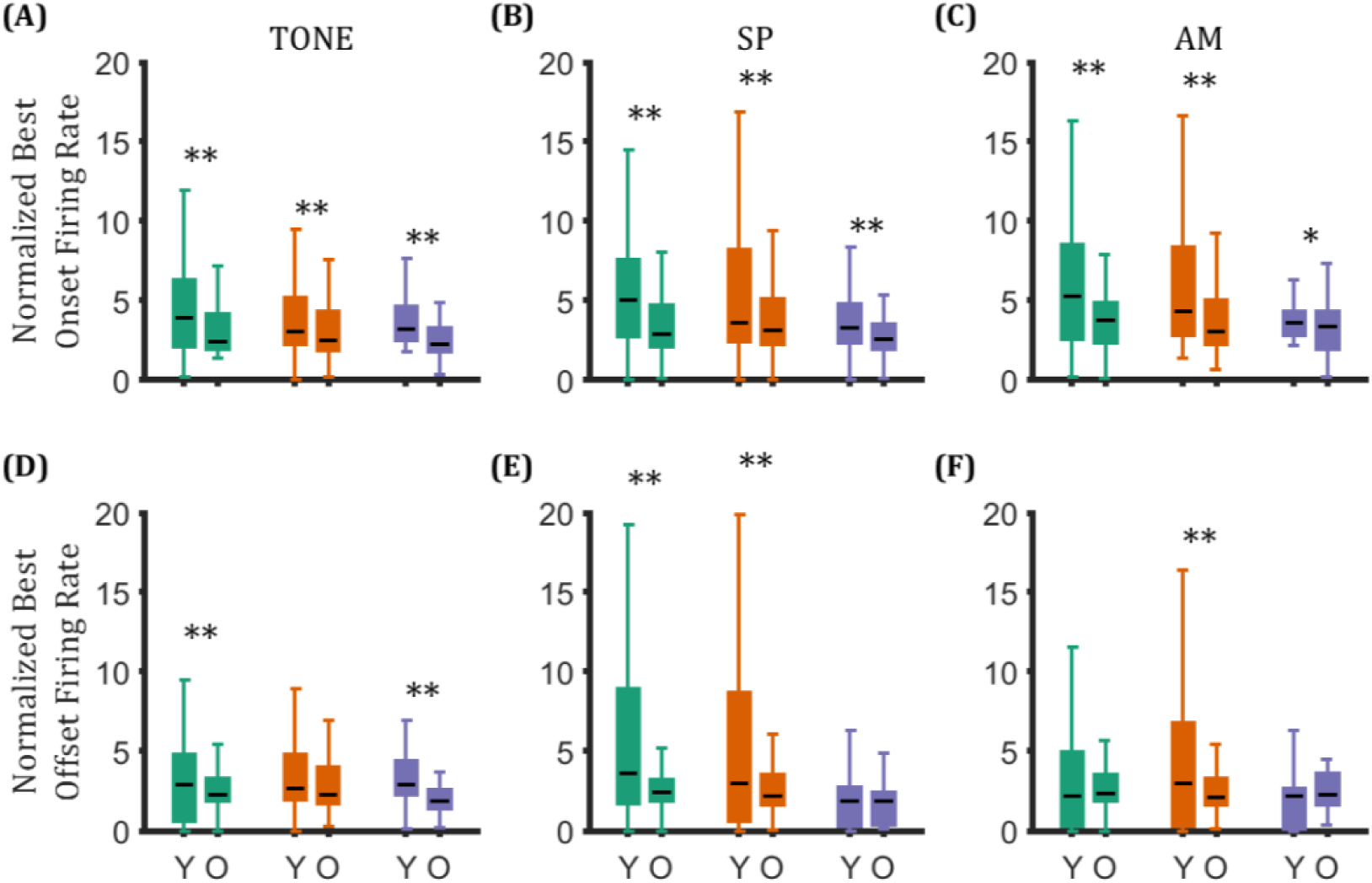
Comparison of the normalized best onset and best offset firing rates between the young and aged monkeys. Box plots illustrate the normalized onset (A-C) and offset (D-F) firing rates across three cortical areas between Monkey Y and Monkey O (**: p<0.01; *: p<0.05; two-sample K-S test).

**Table 4.** Statistical details of the normalized firing rates for ON and OFF responses. Table 4. The median value of the normalized firing rate for each of the three stimulus types (sets of rows) for each of the three cortical areas are shown in column 3-5, and the upper whisker are shown in column 6-8. Statistical p-values (far right column) that are < 0.01 are highlighted in yellow and 0.01 < p <0.05 are in green (two-sample K-S test). There were consistent differences between the young and old monkey across comparisons.

|  | Monkey | Median Value |  |  | Upper Whisker |  |  | Between Monkeys |
| --- | --- | --- | --- | --- | --- | --- | --- | --- |
|  |  | R | A1 | CL | R | A1 | CL |  |
| Norm. FR Best ON (TONE) | Y | 3.92 | 3.08 | 3.19 | 12.57 | 9.53 | 7.88 | R: Y>O, $p=2.14e-3$<br>A1: Y>O, $p=7.66e-4$<br>CL: Y>O, $p=8.11e-6$ |
|  | O | 2.42 | 2.50 | 2.29 | 7.45 | 7.97 | 5.55 |  |
| Norm. FR Best OFF (TONE) | Y | 2.87 | 2.67 | 2.87 | 11.02 | 9.09 | 7.71 | R: Y>O, $p=3.00e-3$<br>A1: Y~O, $p=0.0628$<br>CL: Y>O, $p=6.35e-5$ |
|  | O | 2.27 | 2.28 | 1.84 | 5.47 | 7.49 | 4.27 |  |
| Norm. FR Best ON (SP) | Y | 5.06 | 3.61 | 3.30 | 14.93 | 17.49 | 8.50 | R: Y>O, $p=3.96e-9$<br>A1: Y>O, $p=3.29e-7$<br>CL: Y>O, $p=8.17e-4$ |
|  | O | 2.89 | 3.11 | 2.58 | 8.71 | 9.42 | 6.02 |  |
| Norm. FR Best OFF (SP) | Y | 3.60 | 3.00 | 1.83 | 19.83 | 24.00 | 6.33 | R: Y>O, $p=3.11e-8$<br>A1: Y>O, $p=2.01e-6$<br>CL: Y~O, $p=0.502$ |
|  | O | 2.39 | 2.18 | 1.85 | 5.31 | 6.29 | 4.90 |  |
| Norm. FR Best ON (AM) | Y | 5.30 | 4.29 | 3.57 | 17.51 | 16.97 | 6.65 | R: Y>O, $p=1.64e-5$<br>A1: Y>O, $p=2.72e-6$<br>CL: Y>O, $p=0.0224$ |
|  | O | 3.79 | 3.07 | 3.34 | 8.65 | 9.27 | 7.91 |  |
| Norm. FR Best OFF (AM) | Y | 2.19 | 2.96 | 2.18 | 12.04 | 18.03 | 6.33 | R: Y~O, $p=0.0556$<br>A1: Y>O, $p=1.96e-4$<br>CL: Y~O, $p=0.112$ |
|  | O | 2.38 | 2.13 | 2.26 | 5.92 | 5.70 | 6.70 |  |

We next examined the differences in the onset and offset firing evoked by the best stimulus in individual auditory cortex neurons (Fig. 6; statistical details in Table 5), using raw firing rate data for comparison. As before, the Shapiro-Wilk test indicated that the data were non-parametric, so we applied the Wilcoxon signed-rank test to assess differences between datasets. For area R neurons, the firing rates for onset and offset were not significantly different except for the AM task in the young monkey (Fig. 6A, 6D, and 6G).

**Figure 6.**
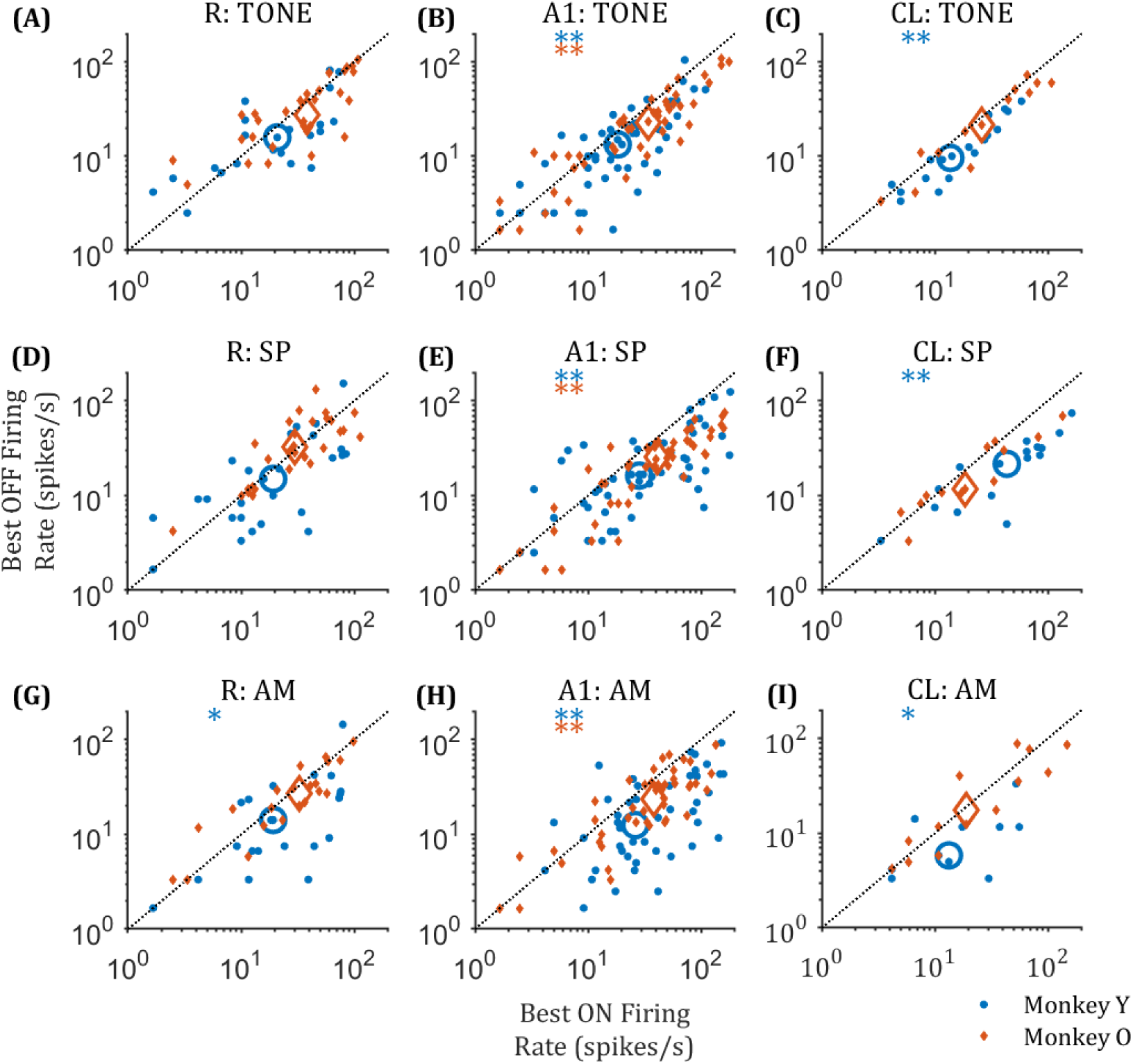
Comparison of the best onset and best offset responses. The black dashed lines show the unity line (no difference). A-C: onset and offset responses evoked by their best tone stimuli. D-F: onset and offset responses evoked by their best location stimuli. G-I: onset and offset responses evoked by their best AM stimuli. The open orange and blue symbols indicate the median value of the onset and offset responses of Monkey Y and Monkey O, respectively (**: p<0.01; *: p<0.05; Wilcoxon signed-rank test).

**Table 5.**
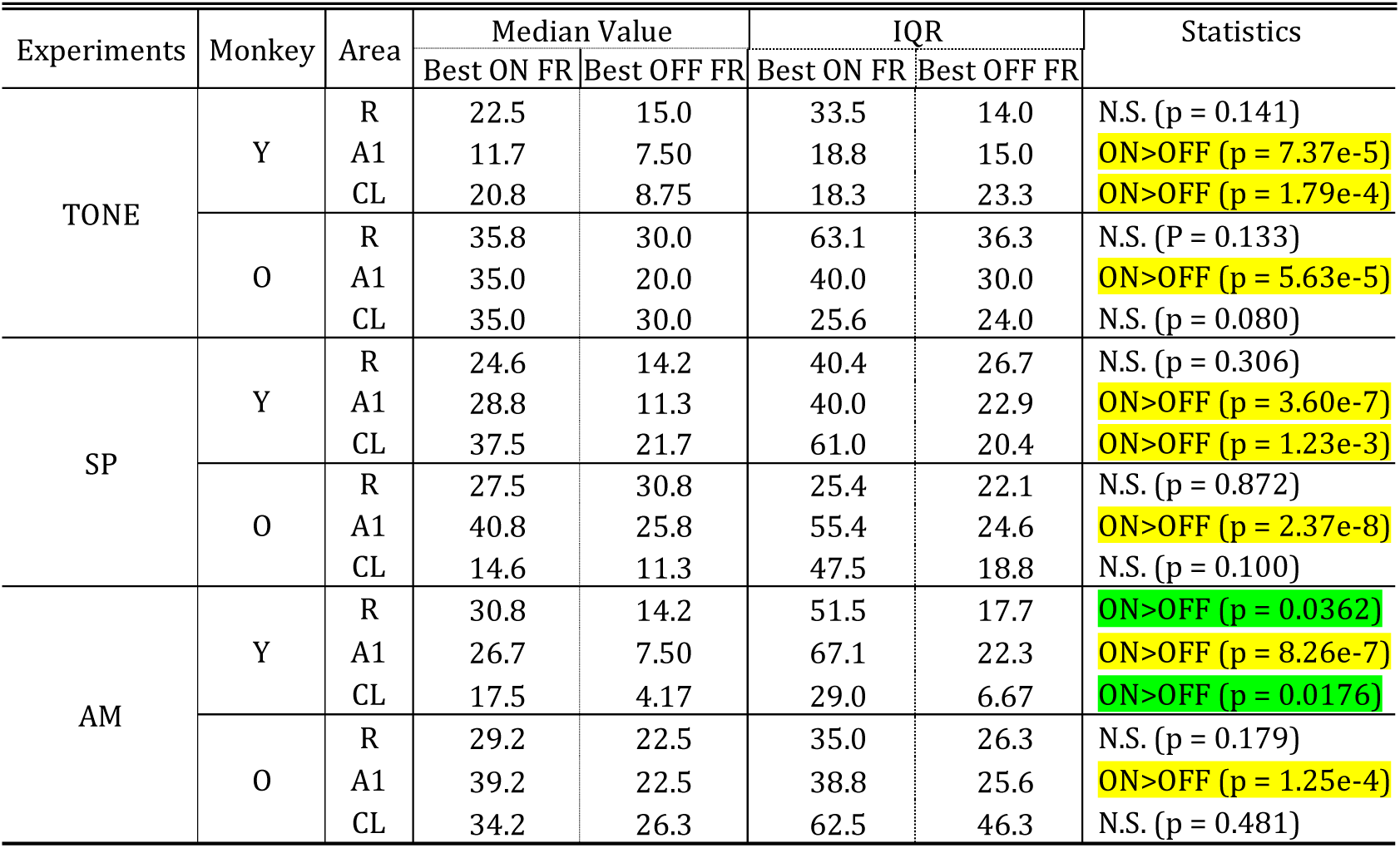
Summary data for Figure 6. Table 5. Statistical significance was tested using the Wilcoxon signed-rank test, with p-values < 0.01 highlighted in yellow, and 0.01<p<0.05 highlighted in green.

| Experiments | Monkey | Area | Median Value |  | IQR |  | Statistics |
| --- | --- | --- | --- | --- | --- | --- | --- |
|  |  |  | Best ON FR | Best OFF FR | Best ON FR | Best OFF FR |  |
| TONE | Y | R | 22.5 | 15.0 | 33.5 | 14.0 | N.S. (p = 0.141) |
|  |  | A1 | 11.7 | 7.50 | 18.8 | 15.0 | ON>OFF (p = 7.37e-5) |
|  |  | CL | 20.8 | 8.75 | 18.3 | 23.3 | ON>OFF (p = 1.79e-4) |
|  | O | R | 35.8 | 30.0 | 63.1 | 36.3 | N.S. (P = 0.133) |
|  |  | A1 | 35.0 | 20.0 | 40.0 | 30.0 | ON>OFF (p = 5.63e-5) |
|  |  | CL | 35.0 | 30.0 | 25.6 | 24.0 | N.S. (p = 0.080) |
| SP | Y | R | 24.6 | 14.2 | 40.4 | 26.7 | N.S. (p = 0.306) |
|  |  | A1 | 28.8 | 11.3 | 40.0 | 22.9 | ON>OFF (p = 3.60e-7) |
|  |  | CL | 37.5 | 21.7 | 61.0 | 20.4 | ON>OFF (p = 1.23e-3) |
|  | O | R | 27.5 | 30.8 | 25.4 | 22.1 | N.S. (p = 0.872) |
|  |  | A1 | 40.8 | 25.8 | 55.4 | 24.6 | ON>OFF (p = 2.37e-8) |
|  |  | CL | 14.6 | 11.3 | 47.5 | 18.8 | N.S. (p = 0.100) |
| AM | Y | R | 30.8 | 14.2 | 51.5 | 17.7 | ON>OFF (p = 0.0362) |
|  |  | A1 | 26.7 | 7.50 | 67.1 | 22.3 | ON>OFF (p = 8.26e-7) |
|  |  | CL | 17.5 | 4.17 | 29.0 | 6.67 | ON>OFF (p = 0.0176) |
|  | O | R | 29.2 | 22.5 | 35.0 | 26.3 | N.S. (p = 0.179) |
|  |  | A1 | 39.2 | 22.5 | 38.8 | 25.6 | ON>OFF (p = 1.25e-4) |
|  |  | CL | 34.2 | 26.3 | 62.5 | 46.3 | N.S. (p = 0.481) |

In stark contrast to area R neurons, we found that the onset responses in A1 had a consistently greater firing rate than the off response across all experiments in both monkeys (Fig. 6B, 6E, and 6H).

Similarly, CL neurons showed that the onset response was greater than the offset response across all three experiments for the young monkey, but not for any experiments in the old monkey (Fig. 6C, 6F, and 6I). Thus, while onset responses were consistently greater than offset responses in both monkeys early in the processing pathway (area A1), this was no longer the case for belt area CL in the old monkey even though it was retained in the young monkey, and in core area R neither neurons from the young or old monkey showed this behavior at the statistically significant level.

### 3. Difference between best onset and offset stimuli

We calculated the best stimulus difference (BSD) in frequency or spatial location between the best onset and offset responses of the neurons for the three types of stimuli, defined as the absolute difference in the best stimulus for onset and offset responses (see Methods). In this part of the data analysis, we applied the Shapiro-Wilk test, and the

Wilcoxon signed rank test to assess the differences between the datasets. The results show that in the SP and AM experiments, there were no significant differences in the BSD values between the three areas within each monkey, nor were there substantial differences when comparing the same areas between the two monkeys (Fig. 7; detailed statistical results in Table 6). The only differences that we saw were in the TONE experiment where CL neurons in the old monkey had a greater difference than both area A1 and area R neurons (Fig. 7A, right panel). This translated into a difference between the two monkeys as BSD (Fig. 7D).

**Figure 7.**
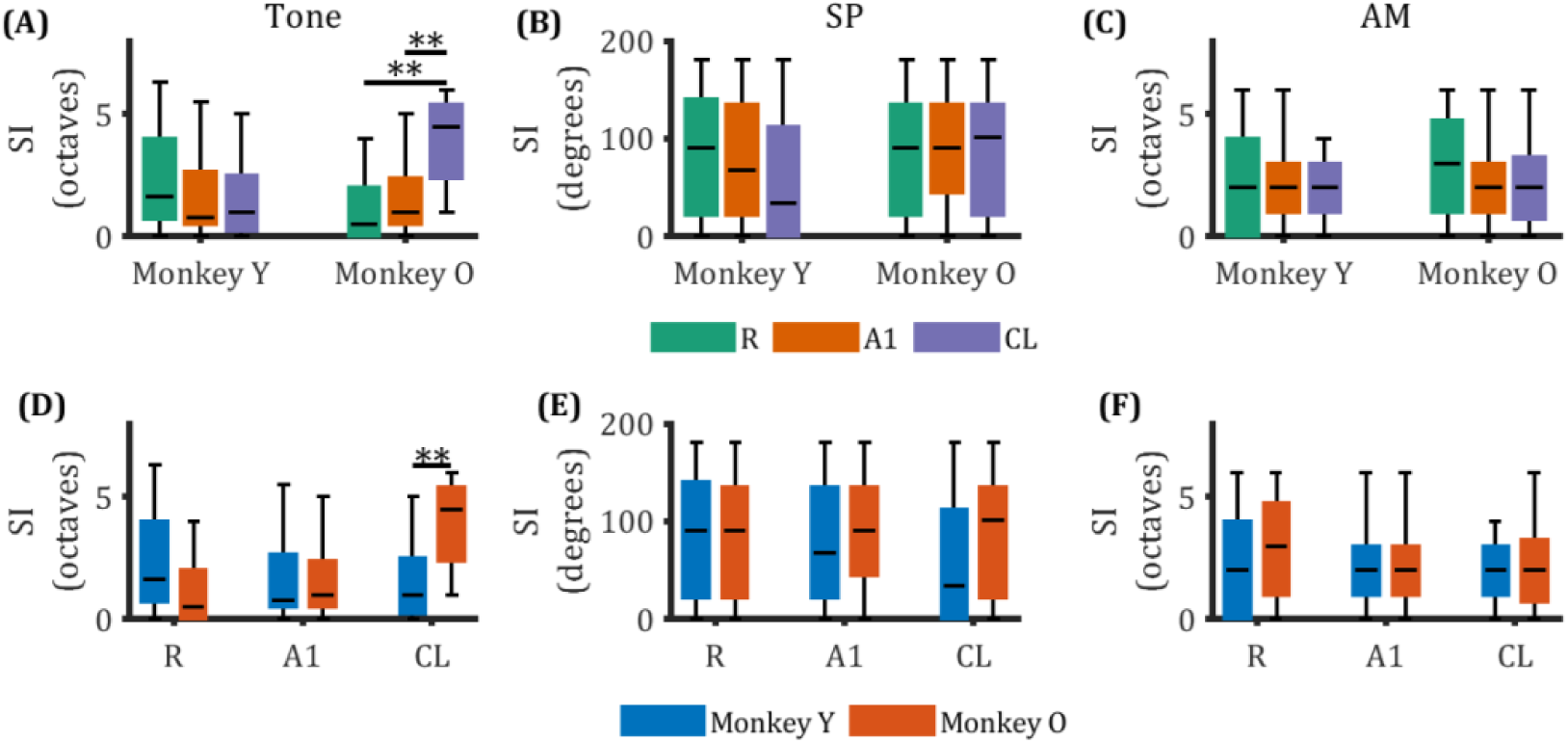
Absolute difference in best onset and offset stimulus (**: p<0.01; *: p<0.05; Wilcoxon signed-rank test).

**Table 6.** Detailed statistical results for the data shown in Figure 7. Table 6. Statistical significance was tested using the Wilcoxon signed-rank test, with p-values < 0.01 highlighted in yellow.

| Experiments | Monkey | Median Value |  |  | Statistics |  |
| --- | --- | --- | --- | --- | --- | --- |
|  |  | R | A1 | CL |  |  |
| TONE | Y | 1.65 | 0.79 | 1.00 | R≈A1: $p=0.146$<br>R≈CL: $p=0.491$<br>A1≈CL: $p=0.757$ | R: Y≈O, $p=0.209$<br>A1: Y≈O, $p=0.989$<br>CL: Y<O, $p=7.86e-3$ |
| | O | 0.50 | 1.00 | 4.50 | R≈A1: $p=0.523$<br>R<CL: $p=3.85e-3$<br>A1<CL: $p=5.37e-3$ | |
| SP | Y | 90 | 67.5 | 33.8 | R≈A1: $p=0.760$<br>R≈CL: $p=0.0892$<br>A1≈CL: $p=0.0573$ | R: Y≈O, $p=0.916$<br>A1: Y≈O, $p=0.429$<br>CL: Y≈O, $p=0.408$ |
| | O | 90.0 | 90 | 101.25 | R≈A1: $p=0.825$<br>R≈CL: $p=0.700$<br>A1≈CL: $p=0.635$ | |
| AM | Y | 2.00 | 2.00 | 2.00 | R≈A1: $p=0.989$<br>R≈CL: $p=0.729$<br>A1≈CL: $p=0.996$ | R: Y≈O, $p=0.860$<br>A1: Y≈O, $p=0.980$<br>CL: Y≈O, $p=1.00$ |
| | O | 3.00 | 2.00 | 2.00 | R≈A1: $p=0.561$<br>R≈CL: $p=0.958$<br>A1≈CL: $p=0.995$ | |

Thus, while there were consistent differences between the magnitude of the onset and offset responses, there were only minor differences in the stimuli that drove those responses.

### 4. Tuning properties of the onset and offset responses

Following our examination of neuronal firing characteristics, we further explored their tuning properties. We measured the spectral, spatial, and temporal tuning bandwidths across the three auditory cortex areas. For this analysis, we applied the Shapiro-Wilk test and the two-sample Kolmogorov-Smirnov test to assess differences between datasets.

Figure 8 presents examples of the tuning functions for tone stimuli, highlighting the onset response and offset responses of two individual area A1 neurons of the young monkey. The raster plots are shown in Fig. 8A and 8C where there are clear onset responses at low frequencies in both neurons. Fig. 8B illustrates the tuning functions from the example neuron shown in Fig. 8A that had highly overlapping responses between its best onset and offset responses, although the offset response was lower, as described for most neurons above. Fig. 8D shows the tuning function from the example of a neuron shown in Fig. 8C with non-overlapping frequency responses between the onset and offset across frequencies.

**Figure 8.**
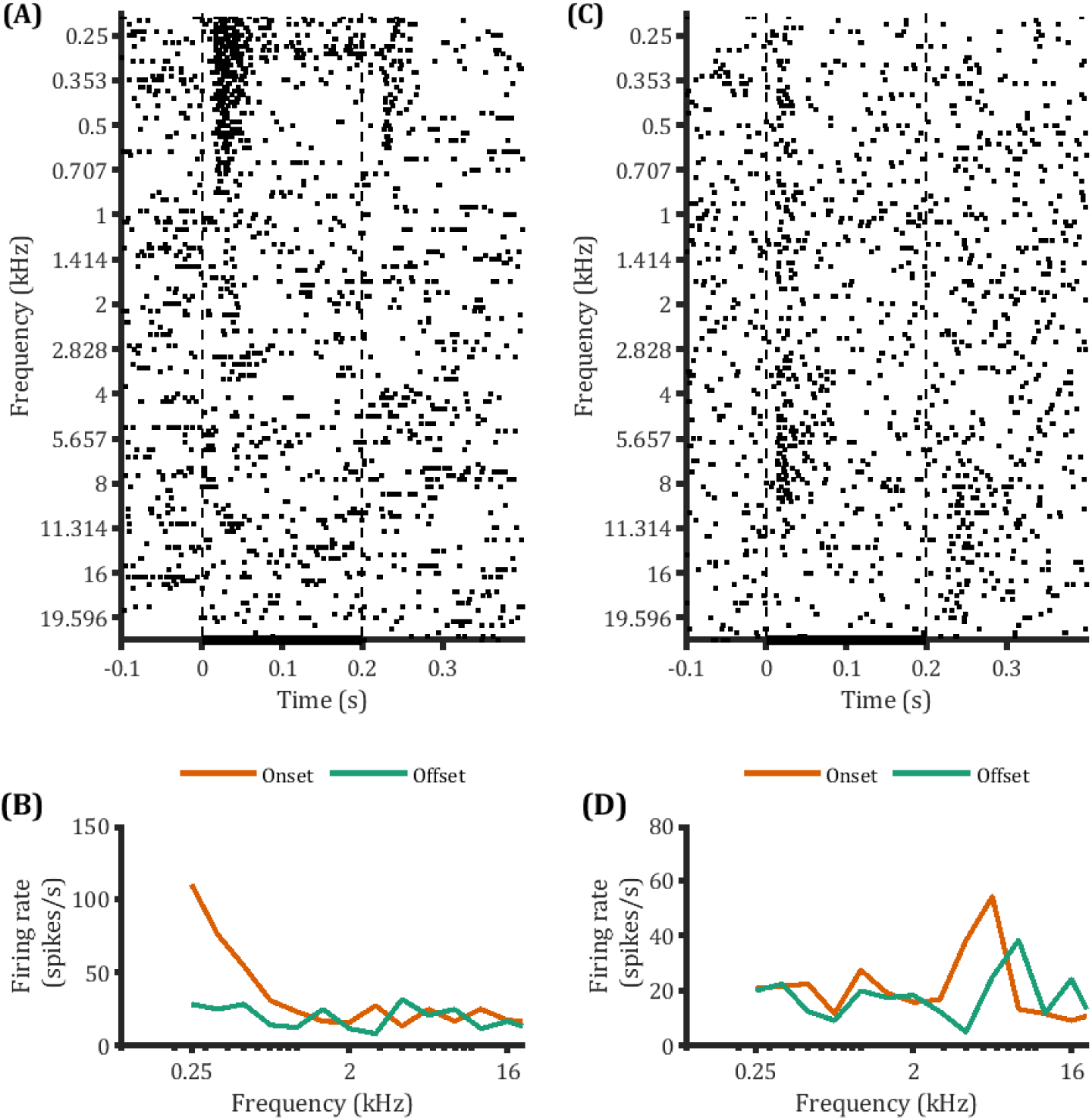
Representative examples of overlapping and non-overlapping A1 cells from the young monkey. A: raster plot of an overlapping cell. B: raster plot of a non-overlapping cell. C: the frequency tuning curves of the onset and offset responses of the cell shown in A. D: the frequency tuning curves of the onset and offset responses of the cell shown in B.

We quantified these impressions across the dataset between the young and old monkey for each cortical area in the TONE experiment (Fig. 9; detailed statistical results in Table 7). We found that there was no significant difference in BWs for onset (Fig. 9A) or offset (Fig. 9B) responses among the three areas in the young monkey. However, in the aged monkey, area R and A1 neurons had significantly narrower onset BWs than area CL neurons (Fig. 9D and 9G), and area R neurons had narrower offset BWs than area CL neurons (Fig. 9D and 9H).

**Figure 9.**
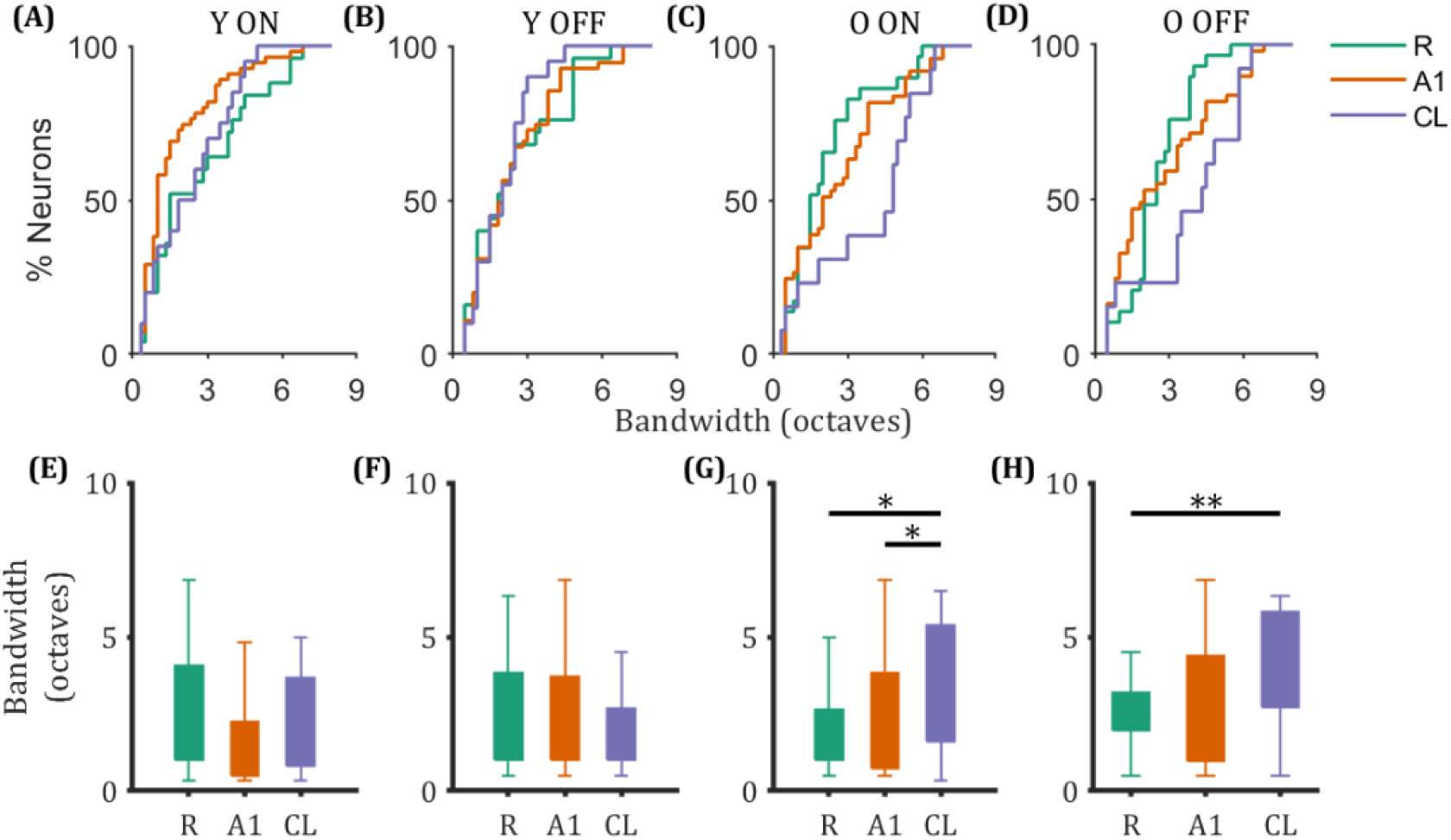
Spectral tuning characteristics of onset and offset in the receptive field subregions of R, A1, and CL neurons in Monkey Y and Monkey O. A-D: the distribution of the neurons’ bandwidths for R, A1, and CL of monkey Y ON (A), monkey Y OFF (B), monkey O ON (C), and monkey O OFF (D). E-H: the comparison of the bandwidths between R, A1, and CL for monkey Y ON (E), monkey Y OFF (F), monkey O ON (G), and monkey O OFF (H) (**: p<0.01; *: p<0.05; two-sample K-S test).

**Table 7.** Statistical details of the analysis shown in Figures 9 and 10 for the TONE experiment. Table 7. Statistical significance was tested using the two-sample K-S test, with p-values < 0.01 highlighted in yellow, and 0.01<p<0.05 highlighted in green.

|  | Monkey | Median Value |  |  | Among Areas | Between Monkeys |
| --- | --- | --- | --- | --- | --- | --- |
|  |  | R | A1 | CL |  |  |
| BW ON (TONE) | Y | 1.50 | 1.00 | 2.17 | R ≈ A1: $p=0.120$<br>R ≈ CL: $p=0.915$<br>A1 ≈ CL: $p=0.137$ | R:<br>Y ON ≈ Y OFF: $p=0.494$<br>O ON ≈ O OFF: $p=0.245$<br>Y ON ≈ O ON: $p=0.470$<br>Y OFF ≈ O OFF: $p=0.209$ |
| | O | 1.50 | 2.00 | 4.83 | R ≈ A1: $p=0.371$<br>R < CL: $p=0.0215$<br>A1 < CL: $p=0.0298$ | A1:<br>Y ON < Y OFF: $p=9.71e-5$<br>O ON ≈ O OFF: $p=0.541$<br>Y ON < O ON: $p=7.58e-3$<br>Y OFF ≈ O OFF: $p=0.392$ |
| BW OFF (TONE) | Y | 1.83 | 2.00 | 2.00 | R ≈ A1: $p=0.681$<br>R ≈ CL: $p=0.486$<br>A1 ≈ CL: $p=0.732$ | CL:<br>Y ON ≈ Y OFF: $p=0.586$<br>O ON ≈ O OFF: $p=0.718$<br>Y ON < O ON: $p=0.0187$<br>Y OFF < O OFF: $p=7.56e-4$ |
| | O | 2.50 | 2.00 | 4.33 | R ≈ A1: $p=0.136$<br>R < CL: $p=7.86e-3$<br>A1 ≈ CL: $p=0.105$ | |

We also compared differences between the young and old monkey across cortical areas (Fig. 10; detailed statistical results in Table 7). We found that area R neurons in both monkeys showed no significant difference in BWs between onset and offset responses (Fig.10A and 10D). In A1, the young monkey exhibited wider offset BWs than onset BWs (Fig. 10E, left panel); while the onset BWs in the aged monkey was wider than in the young monkey (Fig. 10E). In CL, there was no significant difference between onset and offset BWs within each monkey, but the aged monkey had wider BWs than the young monkey for both responses (Fig. 10F). Thus, consistent with the preceding analysis (Fig 7), the responses of area CL neurons in the old monkey drove the noted differences in this analysis of spectral responses.

**Figure 10.**
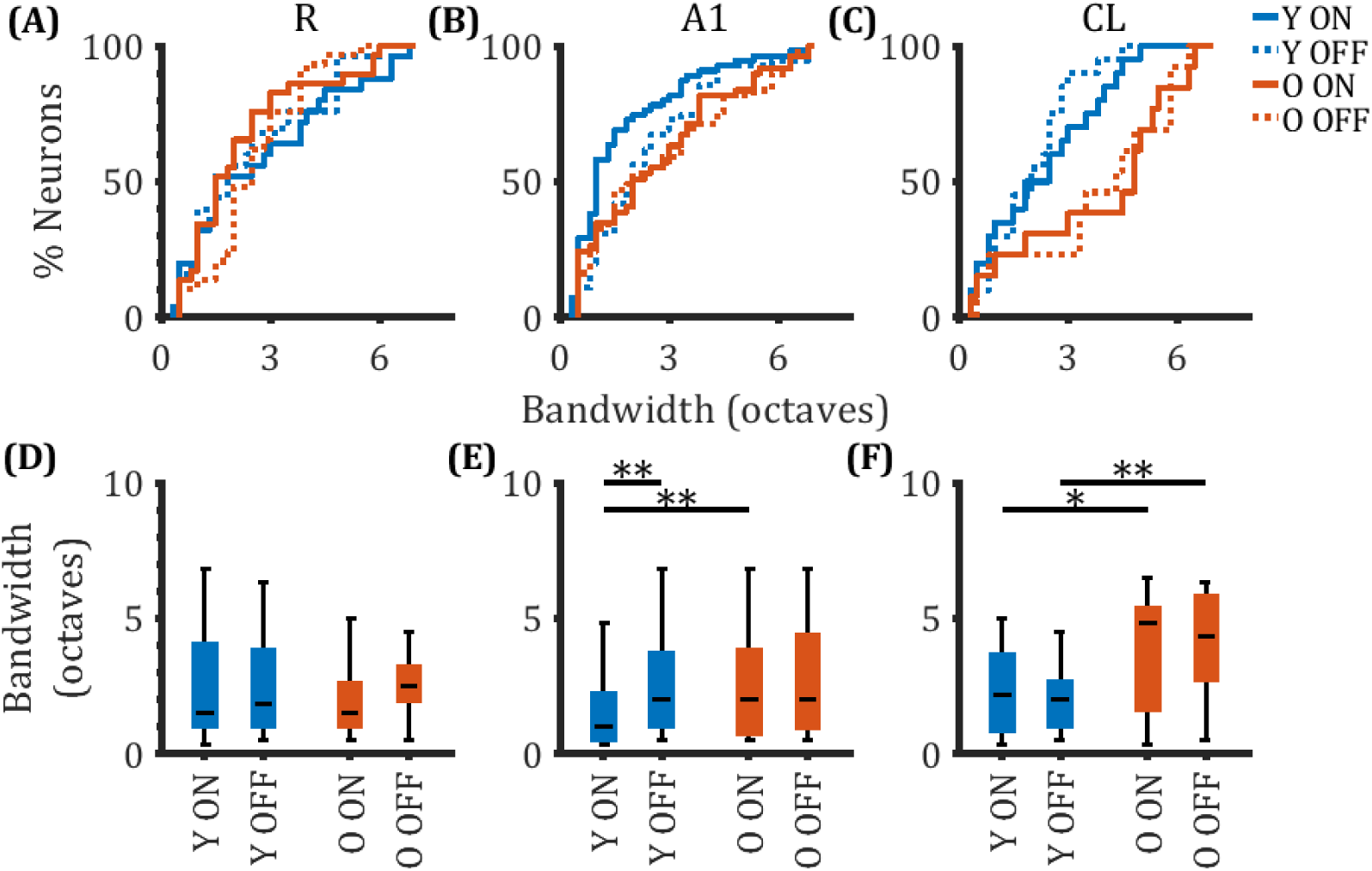
Spectral tuning characteristics of onset and offset in the receptive field subregions of R, A1, and CL neurons in Monkey Y and Monkey O. (**: p<0.01; *: p<0.05; two-sample K-S test).

Applying the same analyses for the SP experiment (Fig. 11 and 12; detailed statistical results in Table 8), we found that the BWs values did not differ significantly among the three areas in the young monkey for either onset or offset responses (Fig. 11A, 11B, 11E, and 11F). However, in the aged monkey, onset BWs in area R neurons were significantly wider than in area CL neurons (Fig. 11C and 11G). In both monkeys, area R neurons exhibited significantly wider onset BWs than offset BWs (Fig. 12A and 12D). Additionally, in area A1 of the young monkey, onset BWs were wider than offset BWs (Fig. 12B and 12E).

**Figure 11.**
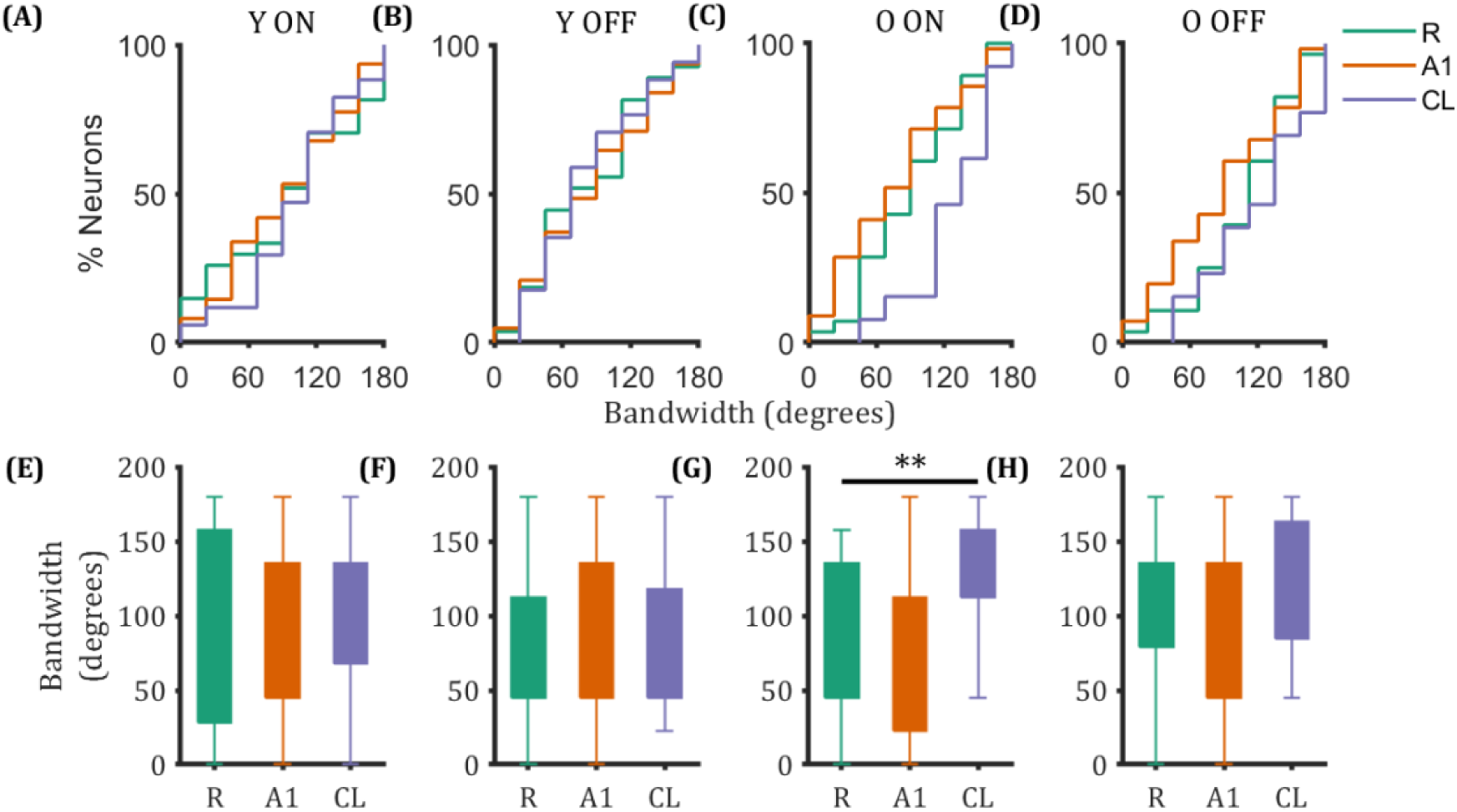
Spatial tuning characteristics of onset and offset in the receptive field subregions of R, A1, and CL neurons in Monkey Y and Monkey O (**: p<0.01; *: p<0.05; two-sample K-S test).

**Figure 12.**
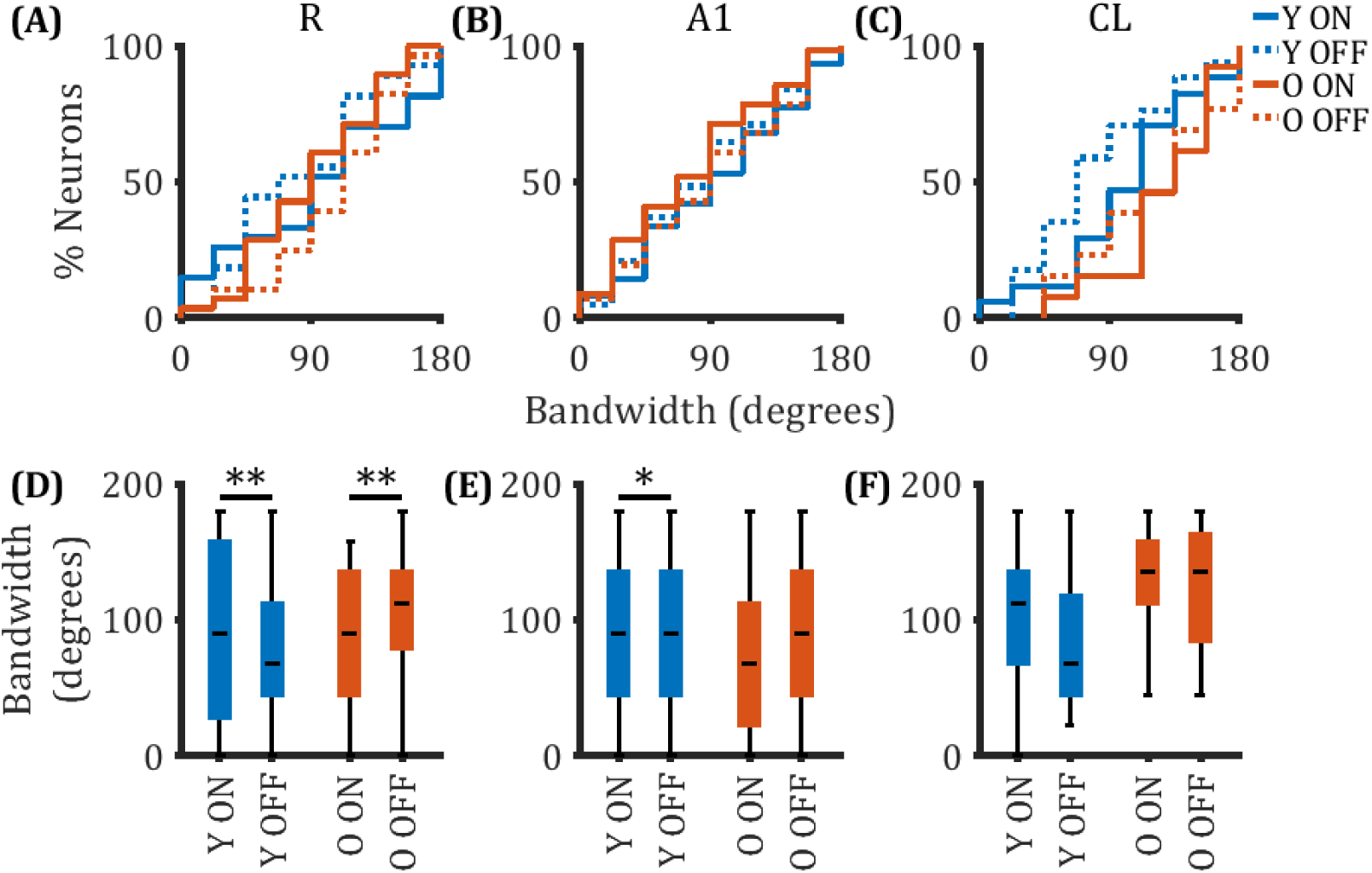
Spatial tuning characteristics of onset and offset in the receptive field subregions of R, A1, and CL neurons in Monkey Y and Monkey O (**: p<0.01; *: p<0.05; two-sample K-S test).

**Table 8.**
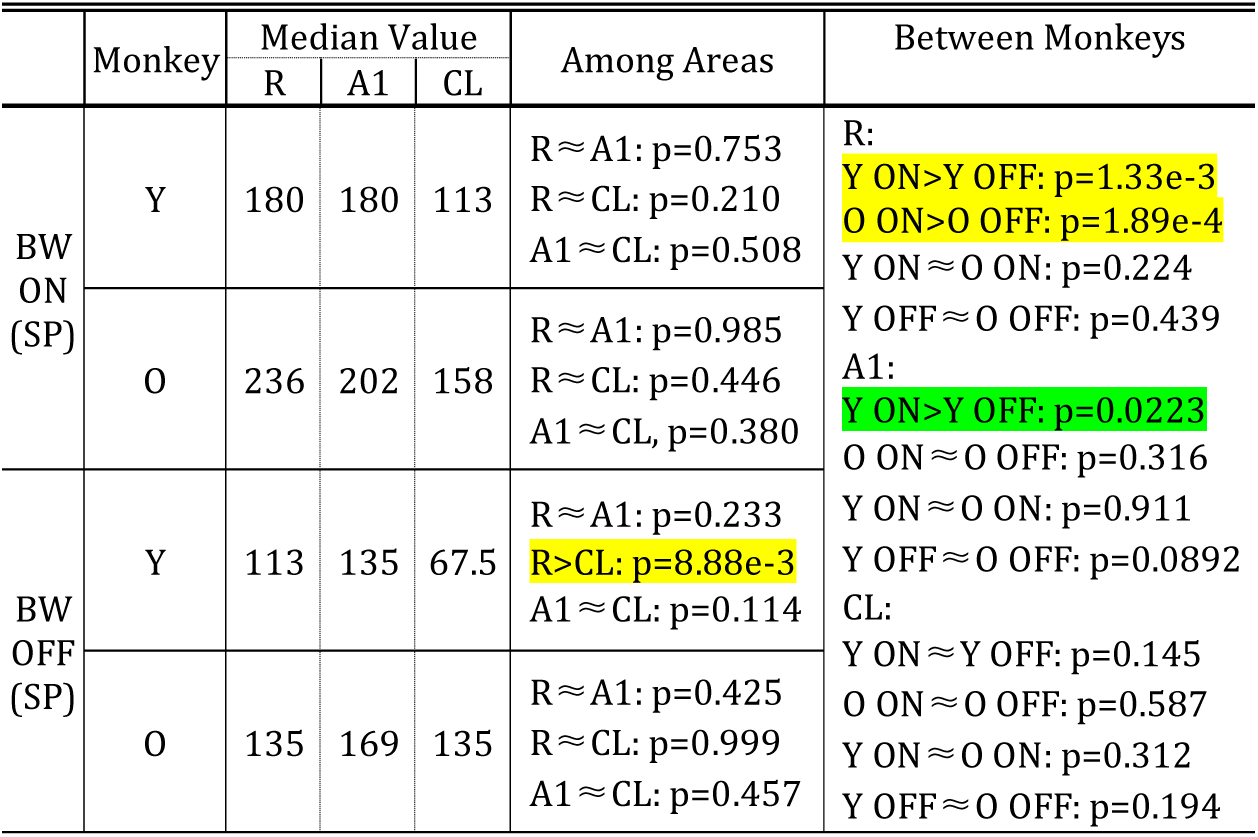
Statistical details for the analysis shown in Figures 11 and 12 on the responses to spatially-varying stimuli. Table 8. Conventions as in Table 7.

Finally, the same analyses for responses during the AM experiment (Fig. 13 and 14; statistical results in Table 9) showed significant BWs differences were observed only between area R and area A1 neurons in the aged monkey (Fig. 13G). Except for CL neurons in the aged monkey, where onset and offset BWs did not differ, all other conditions showed significantly wider onset BWs than offset BWs (Fig. 14D, 14E, and 14F).

**Figure 13.**
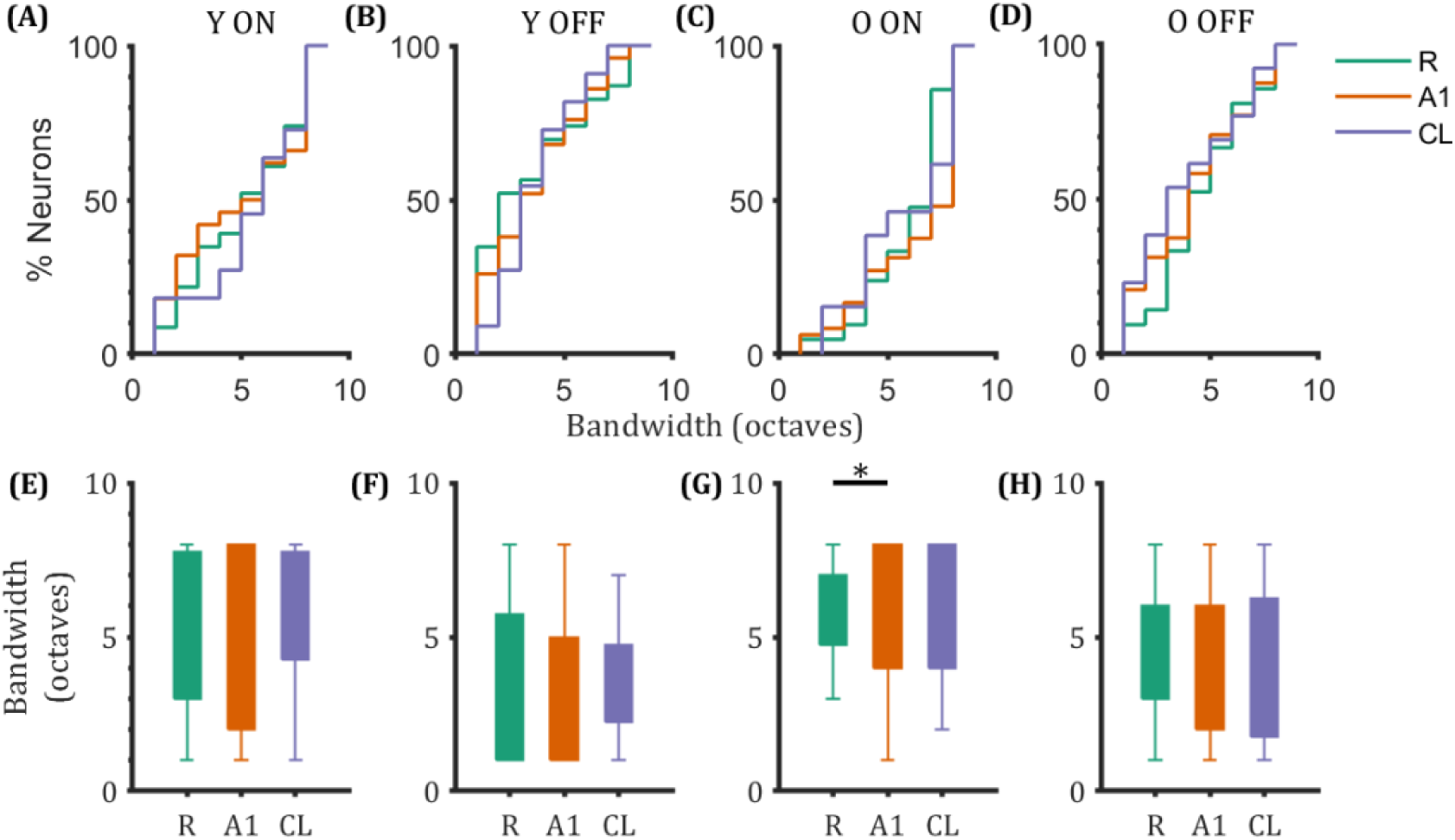
Temporal tuning characteristics of onset and offset in the receptive field subregions of R, A1, and CL neurons in Monkey Y and Monkey O (**: p<0.01; *: p<0.05; two-sample K-S test).

**Figure 14.**
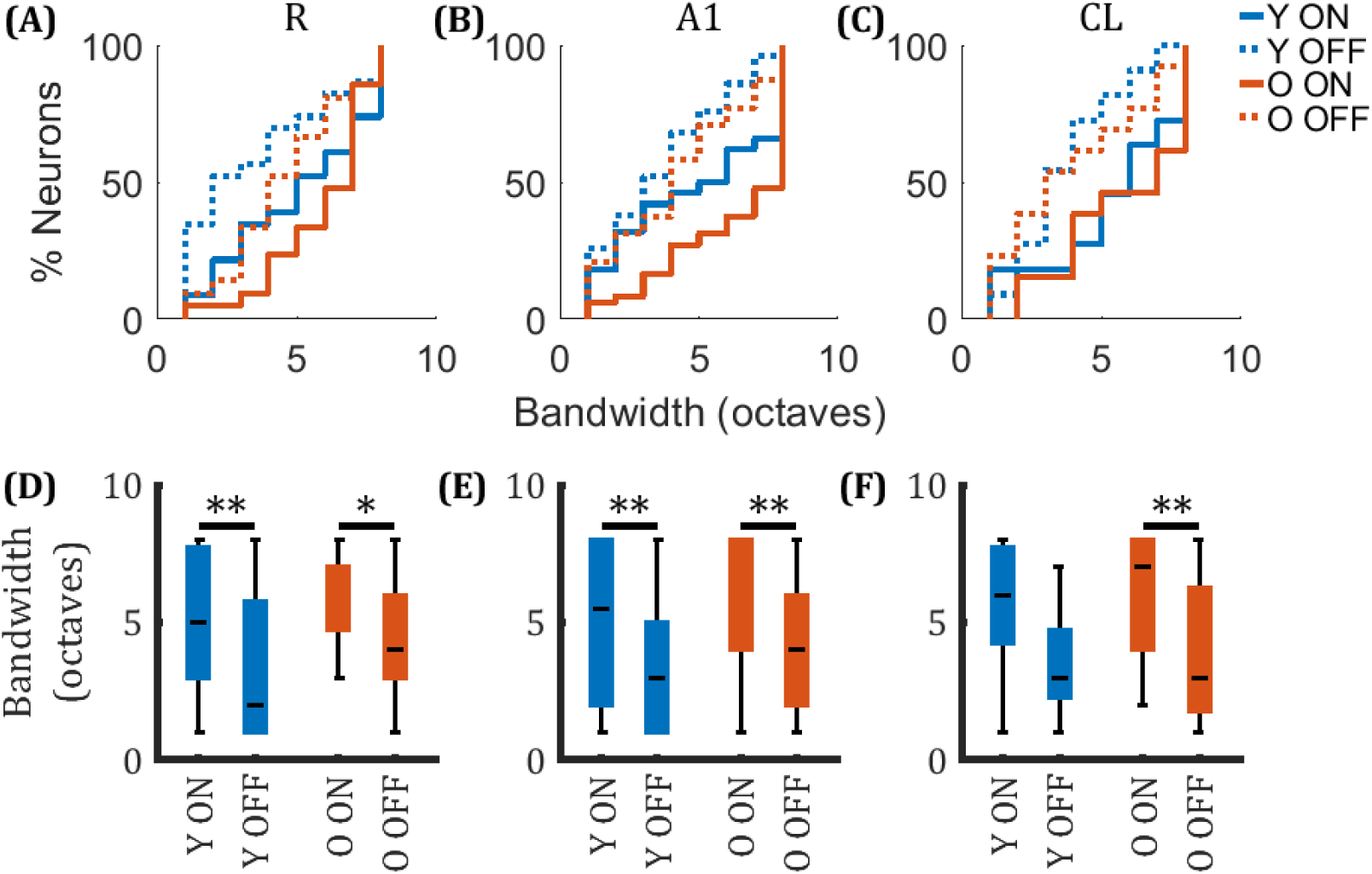
Temporal tuning characteristics of onset and offset in the receptive field subregions of R, A1, and CL neurons in Monkey Y and Monkey O (**: p<0.01; *: p<0.05; two-sample K-S test).

**Table 9.** Statistical details for the analysis shown in Figures 13 and 14 for the AM stimuli. Table 9. Conventions as in Table 7.

|  | Monkey | Median Value |  |  | Among Areas | Between Monkeys |
| --- | --- | --- | --- | --- | --- | --- |
|  |  | R | A1 | CL |  |  |
| BW ON (AM) | Y | 5.00 | 5.50 | 6.00 | R ≈ A1: p=0.994<br>R ≈ CL: p=0.976<br>A1 ≈ CL: p=0.623 | R:<br>Y ON>Y OFF: p=3.23e-3<br>O ON>O OFF: p=0.0187<br>Y ON ≈ O ON: p=0.427<br>Y OFF ≈ O OFF: p=0.0635 |
| | O | 7.00 | 8.00 | 7.00 | R ≈ A1: p=0.885<br>R ≈ CL: p=0.641<br>A1 ≈ CL: p=0.964 | A1:<br>Y ON>Y OFF: p=5.83e-3<br>O ON>O OFF: p=1.68e-6<br>Y ON ≈ O ON: p=0.0718<br>Y OFF ≈ O OFF: p=0.647 |
| BW OFF (AM) | Y | 2.00 | 3.00 | 3.00 | R<A1: p=0.0221<br>R ≈ CL: p=0.672<br>A1 ≈ CL: p=0.964 | CL:<br>Y ON ≈ Y OFF: p=0.0586<br>O ON>O OFF: p=2.93e-3<br>Y ON ≈ O ON: p=0.978<br>Y OFF ≈ O OFF: p=0.999 |
| | O | 4.00 | 4.00 | 3.00 | R ≈ A1: p=0.755<br>R ≈ CL: p=0.672<br>A1 ≈ CL: p=0.925 | |

In summary, across the TONE, SP, and AM experiments, analysis of the BWs revealed distinct age- and region-related differences. In the young monkey, onset and offset BWs were generally comparable across areas A1, R, and CL, with no significant differences among areas. However, in the aged monkey, CL neurons exhibited significantly broader onset and offset BWs than those in areas A1 and R during the TONE experiment. This pattern was reversed in the SP experiment, where area R neurons showed broader onset BWs than area CL neurons. When comparing between monkeys, it is notable that area CL neurons in the aged monkey exhibited significantly expanded BWs in the TONE experiment relative to the young monkey. In contrast, no significant age-related difference was observed in area CL neurons during the SP experiment. In the AM experiment, both monkeys showed significantly broader onset BWs compared to offset BWs across regions, except for the area CL neurons in the young monkey, where no significant difference between onset and offset BWs was found.

## Discussion

Our findings provide information on onset and offset responses in auditory cortical neurons as part of our larger effort to better understand the effects of aging in alert macaque monkey auditory processing (see Recanzone, 2018). In this report we offer a comprehensive examination of how aging influences the responses to the onset and offset of sounds within the core (R and A1) and belt (CL) areas of the auditory cortex. This study provides novel information about the responses between young and aged macaque monkeys passively listening to a variety of different sounds, complimenting previous studies from this and other laboratories (Fishman and Steinschneider, 2009; Tian et al., 2013; Ramamurthy and Recanzone, 2017; Ramamurthy and Recanzone, 2020; Recanzone, 2018).

Recognizing that the sample size is small, with one young and one old monkey contributing to data in this report, it is important to verify that these individuals were not outliers. Our findings in areas A1 and CL are consistent with previous results from our laboratory in different young monkeys (3 individuals) and old monkeys (2 individuals; see Ramamurthy and Recanzone, 2020). We first examined the proportion of neurons exhibiting both onset and offset responses (ON+OFF cells). In previous work from our laboratory using noise burst stimuli, 55.3% of neurons in area A1 of young monkeys and 70.9% in aged monkeys were classified as ON+OFF (Ramamurthy and Recanzone, 2020). In our current SP experiment, which also used noise stimuli, we observed similar or slightly elevated proportions: 69.7% in the young monkey and 73.7% in the aged monkey. For neurons in the caudolateral (CL) belt area, prior results showed ON+OFF proportions of 64.6% in young animals and 59.6% in aged animals. Our current findings revealed 53.1% in the young monkey and 56.5% in the aged monkey, indicating consistency in the relative distributions across studies. We also compared firing rates in the response to each neuron’s best stimulus. Consistent with prior findings in both areas A1 and CL, aged monkeys exhibited higher onset and offset response magnitudes than young monkeys (Ramamurthy and Recanzone, 2020), a pattern that was replicated in the current dataset. Furthermore, spectral tuning bandwidths of onset and offset responses were compared between area A1 and area CL neurons in young monkeys. Previous studies demonstrated that offset responses in A1 tend to have broader spectral tuning than onset responses, whereas neurons in CL showed no significant difference between onset and offset response BWs (Ramamurthy and Recanzone, 2017). Our current findings in the young animal are in agreement with these results. Regarding spatial tuning, earlier work showed that in area A1, the offset tuning BWs were narrower than the onset tuning BWs, while again no significant difference was found between onset and offset tuning in area CL. Our current data similarly reflects these spatial tuning trends. These comparisons strongly indicate that the two monkeys in the current study are consistent with those that we have previously studied and it is very unlikely that these data represent abnormal or outlying idiopathic neural responses.

We should also emphasize that the comparisons are not simply one monkey against the other, but between sets of neurons recorded in the different areas of the two monkeys. Our sample consists of 107 neurons from area R; 249 neurons from area A1; and 104 neurons from area CL. This represents a substantial database for studies of this sort, particularly given the difficulty in obtaining and recording from aged animals (245 neurons total from monkey O). The findings from this study therefore include both confirmations of previously reported phenomena and novel observations that provide additional insights into the onset and offset responses in the normal and aged macaque monkey auditory cortex.

### 1. Altered proportion of ON+OFF neurons between young and aged monkeys

In the young monkey, neurons in the core areas A1 and R exhibited a significantly higher proportion of neurons that responded to both the onset and offset of sounds compared to the neurons in the belt region, across spectral, spatial, and temporally-varying stimuli. This result suggests that in young animals, the core regions are more involved in processing the temporal features of sound, such as their onset and offset, while neurons in the belt region may be engaged in other aspects of sound processing, such as spatial location.

In contrast, however, in the aged monkey this clear distinction between core and belt regions was no longer evident, consistent with previous reports (Juarez-Salinas et al., 2010; Engle et al., 2014; Ng and Recanzone, 2018). The proportion of ON+OFF neurons became more similar across all regions, suggesting that aging reduces the functional specialization between core and belt areas. This loss of specialization may impair the ability of older individuals to process complex sounds, such as speech in noisy environments, where both the onset and offset of sounds are crucial for the appropriate parsing and subsequent processing of the acoustic stream.

When directly comparing the proportion of neurons in the same cortical areas between the two monkeys, we found that only the proportion of ON+OFF neurons in area A1 was significantly lower in the aged monkey compared to the young monkey. This difference was observed only in the spectral and spatial experiments, while no significant difference was found in the temporal experiment. This suggests that aging may particularly affect the processing of spectral and spatial features of sound in area A1, but the temporal processing remains relatively unaffected. Area A1’s role in encoding the frequency and spatial properties of sound may be more susceptible to age-related changes, but further investigation is needed to fully understand the implications for temporal processing.

### 2. Age-related changes in regional firing patterns

After analyzing the differences in the proportion of ON+OFF neurons, we further compared the results from both monkeys in terms of firing rate. Consistent with previous findings (Juarez-Salinas et al., 2010), we observed that the best onset firing rates in all three areas of the aged monkey were significantly higher than those in the young monkey, except for two cases. The same finding was seen for the offset responses (i.e. Fig. 4). These findings are consistent with previous studies in other species, as well as the general change in the excitatory / inhibitory balance observed throughout the ascending auditory neuraxis (Caspary et al., 2008; see Ouda et al., 2015; Gray and Recanzone, 2017). Thus, while less well studied than sub-cortical stations, the neurons in the auditory cortex appear to inherit these changes if not additionally contribute to them.

After comparing the differences in neuronal activity between the two monkeys across different cortical areas, we further examined the firing rate variations within each monkey’s own auditory areas. In the TONE experiment, only the aged monkey showed a significant difference in best offset firing rates, with neurons in area R exhibiting significantly higher firing than those in area A1. Apart from this, no clear regional differences were observed within either monkey. In the AM experiment, significant differences were only found in the young monkey, where the best onset firing rates in the core areas were significantly higher than in CL. No other notable regional differences were observed. However, in the SP experiment, the best onset firing pattern in the young monkey followed a clear trend: R < A1 < CL, indicating a gradual increase in onset firing from the core to the belt region. In contrast, the aged monkey displayed a different pattern—its’ area CL neurons no longer had the highest onset firing rates. Instead, the onset firing rates in area CL neurons were significantly lower than those of neurons in the core areas, consistent with previous findings (Engle and Recanzone, 2013). Examining the best offset firing rates, the young monkey showed minimal differences across cortical areas, except for area A1 neurons, which had significantly higher firing rates than area CL neurons. In contrast, the aged monkey exhibited a distinct offset firing pattern: R > A1 > CL, with area R neurons showing the highest offset activity.

These results suggest that aging alters the hierarchical processing of onset responses, particularly in the belt region. While the young monkey showed distinct areal differences in onset and offset firing under specific conditions, the aged monkey exhibited fewer onset differences but more pronounced offset differences. This shift suggests that aging may diminish the regional specificity of onset responses while amplifying offset-related differences, potentially reflecting compensatory mechanisms or altered neural sensitivity to sound termination.

While increased firing rates were clear in the aged monkey, consistent with previous reports (cited above), we also noted that there was increased spontaneous activity, which has also been seen in auditory cortex of alert primates (Juarez-Salinas et al., 2010). Previous studies have shown that while the overall activity is greater, the tuning properties of auditory cortical neurons in aged monkeys are broadened for spatial (Juarez-Salinas et al., 2010; Engle and Recanzone, 2013), and temporal stimuli (Overton and Recanzone, 2016; Ng and Recanzone, 2018; Ramamurthy and Recanzone, 2020). When we normalized the firing rates to the spontaneous activity we observed a clear pattern: the signal-to-noise ratio of the best onset firing rates were significantly higher for neurons in all cortical areas of the young monkey compared to those of the aged monkey. However, this trend changed for offset firing rates: In the TONE experiment, the offset firing rates of area A1 neurons showed no significant difference between the two monkeys; In the SP experiment, there was no significant difference in the offset firing rates of area CL neurons; In the AM experiment, the offset firing rates of both areas R and CL neurons remained comparable between the two monkeys. These findings indicate that aging differentially affects onset and offset responses, which could clearly impact how older individuals perceive and process ongoing auditory stimuli, or ‘streams’. It remains unclear if the differences due to aging are consequent to the excitatory/inhibitory balance or a shift in processing strategy, as seen previously in AM encoding (Overton and Recanzone, 2016).

Finally, we compared the best onset and offset firing rates across different auditory cortical areas. For both the young and aged monkey, onset responses were essentially the same as offset responses across stimulus types in area R, but consistently higher in the initial cortical processing area (A1), which is diminished at the next core area (R). In contrast, area CL showed a distinct age-related change. While young monkeys exhibited consistently higher onset firing rates than offset firing rates across all experiments, this distinction disappeared in the aged monkey, where onset and offset firing rates became similar. Thus, while core areas retain their functional properties, belt area CL neurons lose this onset-dominant firing pattern, possibly shifting towards an increased reliance on offset responses in aging. This is consistent with amplitude-modulation encoding shifts from a temporal to a rate code in aged monkeys (Overton and Recanzone, 2016). This shift may have implications for how older individuals perceive and process sound transients and is consistent with a reduced ability for fine temporal processing (Ozmeral et al., 2016; Anderson and Karawani, 2020).

### 3. Differences between the best onset and offset tone stimulus characteristics in CL neurons varied between the young and aged monkeys

When comparing the BSD values between the best onset and offset stimuli in these two monkeys in the SP and AM experiments, there were no significant differences in the SI values either between the same monkey’s cortical areas or when comparing the same cortical areas between the young and aged monkeys (Fig. 7). However, in the TONE experiment, the aged monkey’s CL neurons exhibited a significantly larger SIs between the best onset and best offset responses compared to the neurons from the core area.

Furthermore, when comparing the CL neurons between the two monkeys, the difference in SI in the aged monkey was significantly larger than that in the young monkey. This finding suggests that aging may alter the functional relationship between onset and offset responses in area CL, leading to a greater difference in the ON and OFF tuning properties. One possible explanation is that in the young monkey, area CL neurons maintain a more consistent encoding of sound onset and offset, allowing for more integrated processing of sound dynamics. However, in the aged monkey, the increased difference may indicate a functional decoupling between onset and offset responses, potentially reflecting a change in processing strategy away from stricter temporal/spectral encoding toward firing rate encoding (see Overton and Recanzone, 2016).

### 4. Differences in the precision of onset and offset responses

This study also investigated the precision of onset and offset neural responses using a bandwidth (BW) measurement (see Methods). These data are summarized in Figures 9 – 14 and provide a general overview of differences as a function of cortical area and age. The first impression is that, overall, there are few differences that stand out for either cortical area, age, or both. For example, area CL neurons in the old monkey had a broader spectral bandwidth than those in core areas, consistent with previous studies that have measured the entire neural response in anesthetized animals (Rauschecker and Tian, 2004), and old monkey neurons had broader bandwidth than those seen in young monkeys. Smaller changes were noted for spatially-varying stimuli. We did see that young area CL neurons had sharper spatial bandwidths than core areas A1 and R, consistent with previous studies (i.e. Juarez-Salinas et al., 2010; Engle and Recanzone, 2013), although this didn’t reach statistical significance in this study. Similarly, spatial bandwidth was greater in the aged monkey compared to the younger one, again without reaching statistical significance by our criteria. We feel that this is more likely due to the relatively smaller sample size compared to our previous studies as the trends were the same. Finally, for the AM stimuli there was a consistent difference between onset and offset responses, with onset responses being greater than offset responses across comparisons, with all but responses in the young monkey area CL neurons reaching statistical significance.

These results are consistent with previous reports and indicate that both the stimulus type and age can have differential effects on cortical processing, underscoring the importance of taking these parameters into account when making inferences and interpretations of the data. Unfortunately, many studies in alert non-human primates such as this neglect providing the age of the individual animals, and commonly the results of only two individual monkeys are reported and the data are combined as some level of statistical test is applied to show that they are not significantly different. This study has revealed many similarities, but also several differences in temporal encoding of sounds in the alert macaque monkey auditory core (areas A1 and R) and belt (area CL) cortex using three different classes of stimuli, which could have easily been missed depending on the statistical tests (i.e. Herox, 2016).

With the preceding issue in mind, we report all statistical tests and allow the reader to make their own interpretations. We interpret these data to reflect that the functional distinction between core and belt regions was diminished in the aged monkey. In particular, CL neurons in the aged animal exhibited broader spectral tuning and greater divergence between the best frequency for onset and offset responses, indicating a reduced spectral selectivity and temporal precision. Although onset firing rates increased with age, the associated decline in signal-to-noise ratio suggests that this heightened excitability comes at the cost of response fidelity. Such changes may underline age-related challenges in processing spectral, spatial, and temporal aspects of sound that are critical for higher order acoustic processing such as sound discrimination and speech perception, consistent with behavioral observations of deficits in older individuals (Snell et al., 2002; Peelle and Wingfield, 2016). Gaining a deeper understanding of these neural mechanisms offers valuable insights into the decline of auditory perception with age and may inform strategies to alleviate age-related hearing impairments.

## Acknowledgments

The authors would like to thank Brett Bormann, Jennifer Grezlik, Angelly Tovar, Jordan Roberts, Kendall Stewart, Kara Sisson, Melissa Stroud, Katie Neverkovec and Jeffrey Johnson for their help with this project, and Rachel Martin, Adam Cook, Bill Bennett, Rhonda Oates, Kristopher Galang and the California National Primate Research Center for expert animal care.

## Disclaimers

The content of this report is solely the responsibility of the authors and does not necessarily reflect the official views of the University of California

## Author Contributions

DL and GHR conceived and designed the research DL analyzed the data under direction of GHR

DL performed experiments under direction of GHR DL prepared figures with input from GHR

DL prepared the initial versions of the manuscript DL and GHR edited and revised the manuscript

DL and GHR approved the final version of the manuscript

## References

Anderson, S. & Karawani, H. (2020). Objective evidence of temporal processing deficits in older adults. Hear Res 397, 108053. 10.1016/j.heares.2020.108053

Caspary, D.M., Ling, L., Turner, J.G. & Hughes, L.F. (2008). Inhibitory neurotransmission, plasticity and aging in the mammalian central auditory system. J Exp Biol 211(Pt 11), 1781–1791. 10.1242/jeb.013581

David, R.T. & Leathers, C.W. (1985). Behavior and pathology of aging in rhesus monkeys. Am J Phys Anthropol 73(3), 279–414. 10.1002/ajpa.1330730315

Engle, J.R., Gray, D.T., Turner, H., Udell, J.B. & Recanzone, G.H. (2014). Age-related neurochemical changes in the rhesus macaque inferior colliculus. Front Aging Neurosci 6, 73. 10.3389/fnagi.2014.00073

Engle, J.R. & Recanzone, G.H. (2013). Characterizing spatial tuning functions of neurons in the auditory cortex of young and aged monkeys: a new perspective on old data. Front Aging Neurosci 4, 36. 10.3389/fnagi.2012.00036

Engle, J.R., Tinling, S. & Recanzone, G.H. (2013). Age-related hearing loss in rhesus monkeys is correlated with cochlear histopathologies. PLoS One 8(2), e55092. 10.1371/journal.pone.0055092

Fishman, Y.I. & Steinschneider, M. (2009). Temporally dynamic frequency tuning of population responses in monkey primary auditory cortex. Hear Res 254(1-2), 64–76. 10.1016/j.heares.2009.04.010

Frisina, D.R. & Frisina, R.D. (1997). Speech recognition in noise and presbycusis: relations to possible neural mechanisms. Hear Res 106(1-2), 95–104. 10.1016/s03785955(97)00006-3

Gordon-Salant, S. & Fitzgibbons, P.J. (1993). Temporal factors and speech recognition performance in young and elderly listeners. J Speech Hear Res 36(6), 1276–1285. 10.1044/jshr.3606.1276

Gray, D.T., Engle, J.R. & Recanzone, G.H. (2013). Age-related neurochemical changes in the rhesus macaque superior olivary complex. J Comp Neurol 522(3), 573–591. 10.1002/cne.23427

Gray, D.T., Engle, J.R. & Recanzone, G.H. (2014a). Age-related neurochemical changes in the rhesus macaque cochlear nucleus. J Comp Neurol 522(7), 1527–1541. 10.1002/cne.23479

Gray, D.T., Engle, J.R., Rudolph, M.L. & Recanzone, G.H. (2014b). Regional and age-related differences in GAD67 expression of parvalbumin- and calbindin-expressing neurons in the rhesus macaque auditory midbrain and brainstem. J Comp Neurol 522(18), 4074–4084. 10.1002/cne.23659

Gray, D.T. & Recanzone, G.H. (2017). Individual variability in the functional organization of the cerebral cortex across a lifetime: A substrate for evolution across generations. In J. Kaas (Ed.), Evolution of nervous systems 2nd ed., vol. 3, pp. 343–356. Oxford: Elsevier.

Gray, D.T., Rudolph, M.L., Engle, J.R. & Recanzone, G.H. (2013). Parvalbumin increases in the medial and lateral geniculate nuclei of aged rhesus macaques. Front Aging Neurosci., 5, 69. 10.3389/fnagi.2013.00069.

He roux, M. (2016). Inadequate reporting of statistical results. J Neurophysiol., 116(3), 1536–1537. 10.1152/jn.00550.2016.

Juarez-Salinas, D.L., Engle, J.R., Navarro, X.O. & Recanzone, G.H. (2010). Hierarchical and serial processing in the spatial auditory cortical pathway is degraded by natural aging. J Neurosci., 30(44), 14795–14804. 10.1523/JNEUROSCI.3393-10.2010.

Kaas, J.H. & Hackett, T.A. (2000). Subdivisions of auditory cortex and processing streams in primates. Proc Natl Acad Sci U S A, 97(22), 11793–11799. 10.1073/pnas.97.22.11793

Kopp-Scheinpflug, C., Sinclair, J.L. & Linden, J.F. (2018). When sound stops: Offset responses in the auditory system. Trends Neurosci., 41(10), 712–728. 10.1016/j.tins.2018.08.009

Lakatos, P., Pincze, Z., Fu, K.M., Javitt, D.C., Karmos, G. & Schroeder, C.E. (2005). Timing of pure tone and noise-evoked responses in macaque auditory cortex. Neuroreport, 16(9), 933–937. 10.1097/00001756-200506210-00011

Li, H., Wang, J., Liu, G., Xu, J., Huang, W., Song, C., Wang, D., Tao, H.W., Zhang, L.I. & Liang, F. (2021). Phasic Off responses of auditory cortex underlie perception of sound duration. Cell Rep., 35(3), 109003. 10.1016/j.celrep.2021.109003

Livingston, G., Huntley, J., Liu, K. Y., Costafreda, S. G., Selbæk, G., Alladi, S., Ames, D., Banerjee, S., Burns, A., Brayne, C., Fox, N. C., Ferri, C. P., Gitlin, L. N., Howard, R., Kales, H. C., Kivima ki, M., Larson, E. B., Nakasujja, N., Rockwood, K., …, Mukadam, N. (2024). Dementia prevention, intervention, and care: 2024 report of the Lancet standing Commission. Lancet, 404(10452), 572–628. 10.1016/S0140-6736(24)01296-0

Merzenich, M.M. & Brugge, J.F. (1973). Representation of the cochlear partition of the superior temporal plane of the macaque monkey. Brain Res, 50(2), 275–296. 10.1016/0006-8993(73)90731-2

Miller, L.M. & Recanzone, G.H. (2009). Populations of auditory cortical neurons can accurately encode acoustic space across stimulus intensity. Proc Natl Acad Sci U S A, 106(14), 5931–5935. 10.1073/pnas.0901023106

Ng, C.W., Navarro, X., Engle, J.R. & Recanzone, G.H. (2015). Age-related changes of auditory brainstem responses in nonhuman primates. J Neurophysiol, 114(1), 455–467. 10.1152/jn.00663.2014

Ng, C.W. & Recanzone, G.H. (2018). Age-related changes in temporal processing of rapidly-presented sound sequences in the macaque auditory cortex. Cereb Cortex, 28(11), 3775– 3796. 10.1093/cercor/bhx240

Ouda, L., Profant, O. & Syka, J. (2015). Age-related changes in the central auditory system. Cell Tissue Res, 361(1), 337–358. 10.1007/s00441-014-2107-2

Overton, J.A., Cooke, D.F., Goldring, A.B., Lucero, S.A., Weatherford, C. & Recanzone, G.H. (2017). Improved methods for acrylic-free implants in nonhuman primates for neuroscience research. J Neurophysiol, 118(6), 3252–3270. 10.1152/jn.00191.2017

Overton, J.A. & Recanzone, G.H. (2016). Effects of aging on the response of single neurons to amplitude-modulated noise in primary auditory cortex of rhesus macaque. J Neurophysiol, 115(6), 2911–2923. 10.1152/jn.01098.2015

Ozmeral, E.J., Eddins, A.C., Frisina, D.R. Sr. & Eddins, D.A. (2016). Large cross-sectional study of presbycusis reveals rapid progressive decline in auditory temporal acuity. Neurobiol Aging, 43, 72–78. 10.1016/j.neurobiolaging.2015.12.024

Peelle, J.E. & Wingfield, A. (2016). The neural consequences of age-related hearing loss. Trends Neurosci., 39(7), 486–497. 10.1016/j.tins.2016.05.001

Qin, L., Chimoto, S., Sakai, M., Wang, J., & Sato, Y. (2007). Comparison between offset and onset responses of primary auditory cortex ON-OFF neurons in awake cats. J Neurophysiol., 97(5), 3421–3431. 10.1152/jn.00184.2007

Ramamurthy, D.L., & Recanzone, G.H. (2017). Spectral and spatial tuning of onset and offset response functions in auditory cortical fields A1 and CL of rhesus macaques. J Neurophysiol., 117(3), 966–986. 10.1152/jn.00534.2016

Ramamurthy, D.L., & Recanzone, G.H. (2020). Age-related changes in sound onset and offset intensity coding in auditory cortical fields A1 and CL of rhesus macaques. J Neurophysiol., 123(3), 1015–1025. 10.1152/jn.00373.2019

Rauschecker, J.P. (1998). Parallel processing in the auditory cortex of primates. Audiol Neurootol., 3(2-3), 86–103. 10.1159/000013784

Rauschecker, J.P., Tian, B., & Hauser, M. (1995). Processing of complex sounds in the macaque nonprimary auditory cortex. Science, 268(5207), 111–114. 10.1126/science.7701330

Rauschecker, J.P., & Tian, B. (2000). Mechanisms and streams for processing of "what" and "where" in auditory cortex. Proc Natl Acad Sci U S A, 97(22), 11800–11806. 10.1073/pnas.97.22.11800

Rauschecker, J.P., & Tian, B. (2004). Processing of band-passed noise in the lateral auditory belt cortex of the rhesus monkey. J Neurophysiol, 91(6), 2578–2589. 10.1152/jn.00834.2003

Rauschecker, J.P., & Afsahi, R.K. (2023). Anatomy of the auditory cortex then and now. J Comp Neurol, 531(18), 1883–1892. 10.1002/cne.25560

Recanzone, G. (2018). The effects of aging on auditory cortical function. Hear Res, 366, 99–105. 10.1016/j.heares.2018.05.013

Recanzone, G.H. (2000). Response profiles of auditory cortical neurons to tones and noise in behaving macaque monkeys. Hear Res, 150(1-2), 104–118. 10.1016/s0378-5955(00)00194-5

Recanzone, G.H., & Cohen, Y.E. (2010). Serial and parallel processing in the primate auditory cortex revisited. Behav Brain Res, 206(1), 1–7. 10.1016/j.bbr.2009.08.015

Recanzone, G.H., Guard, D.C., & Phan, M.L. (2000a). Frequency and intensity response properties of single neurons in the auditory cortex of the behaving macaque monkey. J Neurophysiol, 83(4), 2315–2331. 10.1152/jn.2000.83.4.2315

Recanzone, G.H., Guard, D.C., Phan, M.L., & Su, T.K. (2000b). Correlation between the activity of single auditory cortical neurons and sound-localization behavior in the macaque monkey. J Neurophysiol, 83(5), 2723–2739. 10.1152/jn.2000.83.5.2723

Snell, K.B., Mapes, F.M., Hickman, E.D., & Frisina, D.R. (2002). Word recognition in competing babble and the effects of age, temporal processing, and absolute sensitivity. J Acoust Soc Am, 112(2), 720–727. 10.1121/1.1487841

Tian, B., Kus mierek, P., & Rauschecker, J.P. (2013). Analogues of simple and complex cells in rhesus monkey auditory cortex. Proc Natl Acad Sci U S A, 110(19), 7892–7897. 10.1073/pnas.1221062110

Tian, B., Reser, D., Durham, A., Kustov, A., & Rauschecker, J.P. (2001). Functional specialization in rhesus monkey auditory cortex. Science, 292(5515), 290–293. 10.1126/science.1058911

World Health Organization. (2021). World report on hearing. https://www.who.int/publications/i/item/9789240020481

Woods, T. M., Lopez, S. E., Long, J. H., Rahman, J. E., & Recanzone, G. H. (2006). Effects of stimulus azimuth and intensity on the single-neuron activity in the auditory cortex of the alert macaque monkey. Journal of Neurophysiology, 96(6), 3323–3337. 10.1152/jn.00392.2006

Yang, W., Zhao, X., Chai, R. & Fan, J. (2023). Progress on mechanisms of age-related hearing loss. Front Neurosci 17, 1253574. 10.3389/fnins.2023.1253574

